# Multi-layered characterization of ∼700,000 conserved noncoding elements within the human genome

**DOI:** 10.64898/2026.06.16.732547

**Authors:** Silvia Fibi-Smetana, Fernando Fernández-Mendoza, Leila Taher

## Abstract

Conserved noncoding elements (CNEs) have been extensively studied for their roles as regulatory elements, particularly enhancers. However, the advent of technologies like ChIP-seq and ATAC-seq has shifted research focus away from comparative genomics. Here, we leveraged data from large-scale projects like ENCODE to address the resulting gap in the comprehensive functional characterization of CNEs.

We first derived a set of ∼700,000 CNEs in the human genome from a 470-way mammalian alignment. Phylogenetic inference identified ∼670,000 conserved elements within primates and ∼240,000 conserved elements across mammals. Our functional genomic analysis revealed that, irrespective of their level of conservation, approximately one third of CNEs exhibit concurrent chromatin accessibility and H3K27 acetylation in at least one of 19 examined tissues and cell lines and thus, are likely to act as *cis*-regulatory elements. Extrapolating these data to additional tissues and cell lines suggested that ∼40% of the CNE repertoire possesses *cis*-regulatory potential. Moreover, we found that the 3D organization of CNEs is non-random; specifically, CNEs are preferentially located toward the centers of topologically associating domains. CNE co-activation networks derived from chromatin accessibility and active histone marks revealed that evolutionary constraints acting on CNEs functioning as *cis*-regulatory elements reflect not only their isolated individual role, but their topological context.

To summarize, we have generated a novel catalog of CNEs annotated with empirical *cis*-regulatory evidence. While evolutionary constraint and regulatory function are clearly linked, a comprehensive understanding of their interplay remains elusive. This resource provides a foundation for exploring this relationship systematically.

**Significance statement:** Previous studies have investigated conserved noncoding element (CNE) evolution, epigenomic landscapes, and 3D genome organization separately, yet a systematic framework integrating these dimensions has been lacking. Here, we identified ∼700,000 CNEs, including ∼240,000 CNEs deeply conserved across mammals, and show that a large fraction display enhancer-associated epigenomic signatures and are preferentially enriched within TAD centers, highlighting their regulatory relevance. By generating and analyzing this CNE catalog within an integrated evolutionary, epigenomic, and 3D genomic context, our study bridges a critical gap and provides a comprehensive resource to better understand regulatory architecture and its potential contribution to disease-associated variation.

## Introduction

Conserved noncoding elements (CNEs) were first observed in the late 1990s and have been characterized multiple times since the beginning of the post-genomics era. A landmark event in the field was the discovery that over half of the 481 ultraconserved elements (UCEs)—defined as segments ≥ 200 bp with 100% identity across human, mouse, and rat—are noncoding (Bejerano et al. 2004). This finding was initially difficult to reconcile with prevailing models of sequence evolution, yet similar patterns were subsequently observed across broader evolutionary distances; for example, whole-genome comparisons between human and pufferfish (*Takifugu rubripes*) identified nearly 1,400 highly conserved noncoding elements, suggesting the existence of noncoding sequences essential across all vertebrates (Woolfe et al. 2005). CNEs are not uniformly distributed throughout the genome, but clustered around developmental genes (Woolfe et al. 2005), or within gene-poor regions, commonly referred to as gene deserts (Nobrega et al. 2003).

Many CNEs have been functionally validated as *cis*-regulatory elements, most notably enhancers, across diverse cellular contexts (Leypold and Speicher 2021). This function was evidenced as early as 1995, when comparing the noncoding genomic sequences of distantly related species such as *Fugu* and mouse emerged as an effective strategy for identifying *cis*-regulatory elements (Aparicio et al. 1995). The principle of evolutionary conservation subsequently served as the foundation for the VISTA Enhancer Browser, a database of tissue-specific developmental enhancers validated using transgenic assays (Pennacchio et al. 2006). More recently, massively parallel reporter assays (MPRAs) have facilitated high-throughput interrogation of CNEs, revealing that nearly half of the human-accelerated regions (HARs) act as enhancers in neural contexts, with the majority of them exhibiting functional differences relative to chimpanzee orthologs (Girskis et al. 2021). Furthermore, MPRAs indicate that ∼800 of ∼10,000 human-specific deletions in CNEs underpin regulatory divergence—typically manifesting by increased regulatory activity (Xue et al. 2023). Beyond their role as *cis*-regulatory elements, CNEs have also been connected to 3D genome organization. Specifically, the borders of CNE clusters have been shown to co-localize with the boundaries of topologically associating domains (TAD), especially those encompassing key developmental genes (Harmston et al. 2017).

The advent of chromatin immunoprecipitation followed by sequencing (ChIP-seq) and high-throughput chromatin accessibility assays has enabled the systematic and comprehensive mapping of regulatory elements across the human genome. Large-scale projects, including ENCODE (ENCODE Project Consortium et al. 2020), FANTOM (Andersson et al. 2014) and Roadmap Epigenomics (Nijim et al. 2025) have generated an unparalleled catalog of regulatory elements across diverse tissues and cell types, greatly facilitating CNE characterization. For instance, a systematic intersection of ∼14,000 UCEs with thousands of ENCODE datasets revealed that these sequences are enriched for SCREEN-predicted candidate *cis*-regulatory elements, transcription factor binding sites, and DNase I hypersensitive sites, with odds ratios exceeding 20 within the embryonic central nervous system tissues (Cummins et al. 2024). Similarly, after screening ∼330,000 CNEs for cetacean-specific sequence divergence, ENCODE ChIP-seq data was used to identify 745 and 1,786 CNEs enriched for H3K27ac and H3K4me1, respectively, which likely function as *cis*-regulatory elements during limb bud initiation stage (Zhang et al. 2025). Furthermore, in GM12878 cells, CNEs identified across ten primate species exhibited enrichment for H3K27ac and H3K79me2 compared to random genomic regions, yet were depleted for DNaseI hypersensitive sites (Hettiarachchi 2022), indicating that while CNEs possess the hallmarks of active or primed enhancers, they are not readily accessible. More generally, while a case study focusing on ChIP-Seq for p300/CBP, H3K4me1, and H3K4me3 and DNaseI hypersensitivity data for two cell types could ascribe regulatory functions to 20,000 of 700,000 unannotated CNEs (Hemberg et al. 2012); by extrapolating this data to additional cell types and conditions, the authors estimated that 80% of CNEs likely function as distal regulatory elements. While these studies provide valuable insight, a systematic investigation integrating CNE evolution, the epigenomic landscape, and 3D genome architecture still remains to be conducted. Here, we present a comprehensive characterization of nearly 700,000 CNEs in the human genome using publicly available data.

## Results

### Evolutionary profiling of the human genome identified over 700,000 CNEs, including ∼240,000 elements deeply conserved across mammals

To bridge the gap between evolutionary constraint and *cis*-regulatory activity, we first established a comprehensive dataset of conserved noncoding elements (CNEs) in the human genome by applying a series of filtering steps to 29 million phastCons elements derived from 470-way alignments (Figure 1a; Methods). Assuming that closely spaced conserved elements may represent single functional units, we initially merged phastCons elements within 50 base pairs of each other if the resulting genomic region overlapped with the original phastCons elements by at least 50% of its length (Methods), obtaining 11.5 million elements. Excluding 10.3 million elements shorter than 50 bp, which were deemed unlikely to be functionally relevant, and 199,527 elements annotated as noncoding RNAs, yielded a final set comprising 191,659 conserved coding elements (CDSs) and 784,915 conserved noncoding elements (CNEs; Figure 1b).

**Figure 1.**
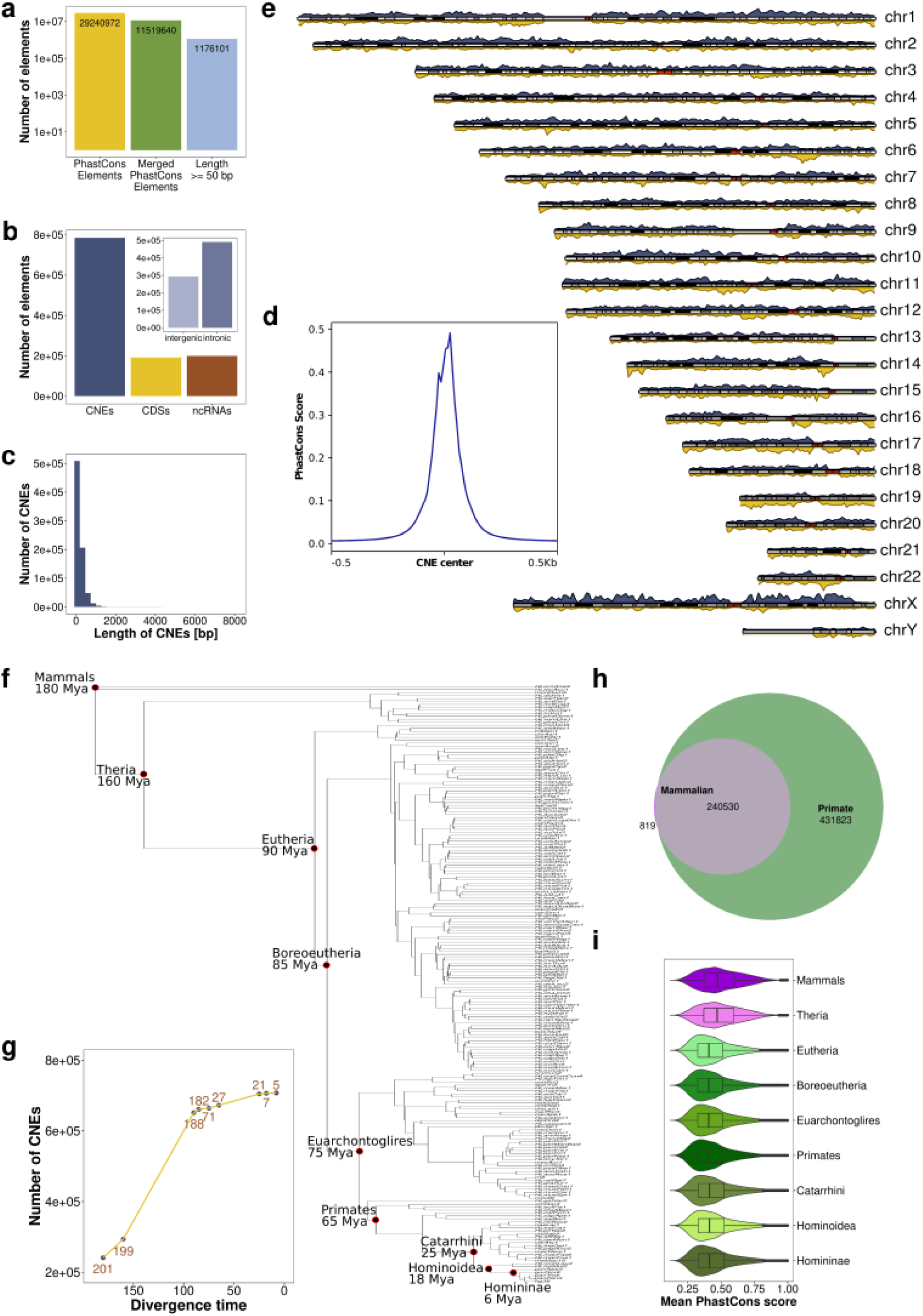
Mammalian CNEs represent a highly constrained core within a larger, more recent repertoire. **a)** Barplot representing the number of conserved elements remaining after each successive filtering stage of the identification pipeline. Starting with 29 million phastCons elements, merging closely spaced ones resulted in 11.5 million elements. Excluding elements shorter than 50 bp yielded 1.2 million conserved elements. **b)** Functional annotation of conserved elements. The main barplot displays how the identified conserved elements are distributed across three categories: coding sequences (CDSs), noncoding RNAs (ncRNAs), and approximately ∼800,000 conserved noncoding elements (CNEs). The inset provides a breakdown of the CNE subset, specifically comparing the number of elements located in intergenic and intronic regions **c)** The histogram displays the length in base pairs (bp) of ∼700,000 CNEs located on human autosomes. The distribution shows a left-skewed profile, indicating a prevalence of shorter elements, with a median of 131 bp. All subsequent panels (d-i) refer to this autosomal dataset. **d)** CNE centers are more conserved than their boundaries based on phastCons score. **e)** Chromosome plot displaying non-uniform distribution of CNEs in grey and genes in yellow in the human genome. **f)** Divergence and ancestral reconstruction of the vertebrate lineage. The phylogenetic tree illustrates the evolutionary relationships among the studied species, with branch lengths representing relative genetic distances. Clades of interest are labeled at their respective ancestral nodes. Divergence times are indicated in millions of years (Mya) for each such node. **g)** Temporal dynamics of CNE emergence across the mammalian lineage. Scatter plot of the number of autosomal CNEs originating at specific ancestral nodes as a function of divergence time (in millions of years, Mya) relative to the human genome. Points denote ancestral nodes of interest within the phylogeny. Brown labels indicate the number of extant species descending from each ancestor. **h)** Intersection between mammalian and primate CNEs displaying that CNEs conserved in the mammalian ancestor are mainly a subset of CNEs conserved in the primate ancestor. **i)** PhastCons scores in noncoding elements conserved in the mammalian and therian ancestor were higher than in elements conserved in younger clades. **ALT TEXT:** Graphs on CNE characterization and evolution, illustrating their distribution in the human genome and evolutionary conservation across the mammalian tree.

CNEs ranged from 51 to 8,006 bp in length, with a median of 131 bp, and represented 5% of the human genome (Figure 1c). As expected, they were on average less strongly conserved than CDSs (mean phastCons scores of 0.44 and 0.65, respectively; Methods), but more strongly conserved than random genomic regions (mean phastCons score of 0.09). In particular, the cores of the CNEs were more conserved than their boundaries (Figure 1d). Our analysis of common human SNPs confirmed that CNEs are also under purifying selection in humans, displaying a markedly lower SNP densities compared to random genomic regions (one-sided Wilcoxon’s test, *P*-value < 2.2×10^−16^). CNEs were distributed across all chromosomes (Figure 1e), with their frequency being proportional to the chromosome length (Pearson’s correlation coefficient R² = 0.92, *P*-value = 1.91×10^−10^), with an average of 32,705 CNEs per chromosome. Most (63%) CNEs were intronic (Figure 1b). The median number of SNPs within both mammalian and primate CNEs was 0. To avoid sex chromosome-related biases, we restricted subsequent analyses to the 709,905 autosomal CNEs.

Next, to reconstruct the evolutionary history of the CNEs, we first identified the orthologs to our human CNEs across the mammalian phylogeny (Figure 1f). For robust phylogenetic inference, we restricted the analysis to the 201 species with high-quality genome assemblies (Methods). As expected, the number of orthologs decreased with increasing phylogenetic distance from humans. For example, we identified ∼707,000 orthologs in chimpanzees, but only ∼259,000 in opossum. Glires deviated from this general trend, yielding fewer orthologs (e.g., ∼550,000 in mice and 606,000 in rabbits) than the more distantly related Carnivora, such as dogs (654,000) and foxes (650,000). This finding aligns well with rodents undergoing accelerated evolutionary rates compared to other mammals, which makes ortholog identification more challenging in this group (Wu and Li 1985). From these data, we then inferred CNE gains and losses throughout the mammalian tree of life (Methods). Thus, consistent with the expectation that CNE numbers generally decrease with increasing phylogenetic depth, we identified 241,349 CNEs as having originated in the reconstructed last common ancestor (LCA) of mammals, the vast majority of which are represented within the 672,353 CNEs inferred in the more recent primate ancestor (Figures 1g and h; Supplementary Table 1). The evolutionary age of the CNEs correlated directly with the degree of evolutionary constraint they exhibit. Specifically, CNEs shared across the mammalian or therian clades consistently display significantly higher phastCons scores compared to those conserved within more recently diverged lineages (Figure 1i). This observation underscores the critical and long-established functional roles of deeply conserved noncoding sequences. CNEs restricted to younger clades are likely to reflect more recently acquired, clade-specific adaptations or functions under less stringent evolutionary constraint, potentially allowing for greater regulatory plasticity.

Building on these insights into CNE evolution, we next sought to link their varying conservation levels to specific functional roles by characterizing their genomic location and regulatory potential.

### Functional profiling revealed that most CNEs act as *cis*-regulatory elements

Approximately 3% of mammalian and 1% of primate CNEs comprised a transcription start site (TSS), distinguishing a minority of CNEs as candidate promoters. The median distance between the mammalian CNEs and their nearest protein-coding gene(s) was 39,460 bp (ranging from 39 bp to 2,473,539 bp), with a total of 13,476 protein-coding genes identified as nearest neighbors (Figure 2a). Primate CNEs exhibited a similar median (32,999 bp) and identical minimum and maximum range, and were found to have 16,340 distinct nearest neighboring protein-coding genes. In both cases, nearest neighbor genes were related to organismal development and cell differentiation (Methods). Processes specifically associated with mammalian CNEs were axonogenesis, Wnt signaling pathway, and eye development. Primate CNEs, on the other hand, were linked to visual system development, ameboidal-type cell migration, and muscle organ development. These findings underscore the fundamental involvement of CNEs in orchestrating developmental processes, with distinct lineage-specific functional roles.

**Figure 2.**
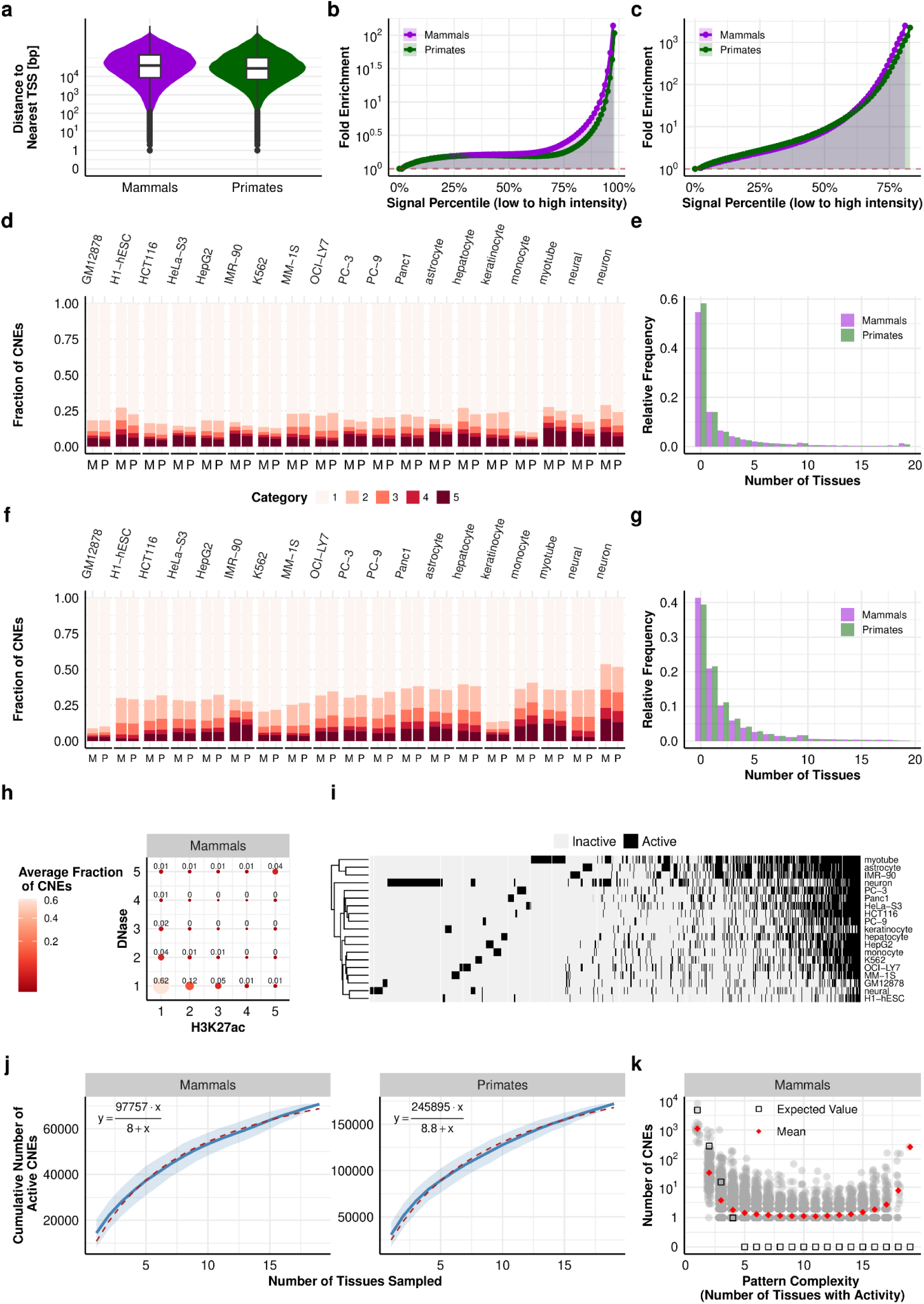
A substantial fraction of CNEs act as *cis*-regulatory elements. **a)** Distance to nearest TSS for mammalian CNEs and primate CNEs. **b)** DNase I hypersensitivity fold-enrichment for mammalian and primate CNEs relative to random genomic regions. Mammalian CNEs featured generally stronger enrichment compared to their primate counterparts. **c)** H3K27ac fold-enrichment for mammalian and primate CNEs relative to random genomic regions. Mammalian CNEs featured generally stronger enrichment compared to their primate counterparts. **d)** Distribution of CNEs by chromatin accessibility. Fractions of mammalian (M) and primate (P) CNEs partitioned into five categories based on their DNase I hypersensitivity (DHS) signal, ranging from Category 1 (closed) to Category 5 (open). **e)** Tissue-specificity of accessible CNEs. Frequency distribution of mammalian and primate CNEs categorized as accessible (Categories 4 and 5) across 19 tissues. A relatively high proportion of CNEs was constitutively accessible in all tissues. **f)** Distribution of CNEs by H3K27ac and **g)** Tissue-specificity of acetylated CNEs. Analogous to panels (d) and (e), but utilizing H3K27ac signal for categorization and tissue-specificity analysis. **h)** Enrichment of CNEs across DNase I and H3K27ac Categories. Bubble size, color intensity, and labels represent the mean fraction of CNEs across all tissues for each combination of DNase I hypersensitivity and H3K27ac categories. The data shows a strong positive correlation between accessibility and acetylation states. See Supplementary Figure 2 for primate data. **i)** Heatmap illustrating the activity profiles of mammalian CNEs across 19 tissues. Active CNEs are those in Category 4 or 5 for both DNase I hypersensitivity and H3K27ac. Only CNEs active in at least one tissue are shown. The distribution reveals a clear bimodal signature, with CNEs predominantly exhibiting either ubiquitous activity or high tissue-specificity. See Supplementary Figure 3 for primate data. **j)** Saturation analysis of CNE activity. Curves estimating the cumulative fraction of mammalian and primate active CNEs as a function of the number of tissues sampled. The solid blue line represents the mean accumulation across 100 random permutations of the 19 tissues under investigation, with the shaded area indicating the standard deviation (SD). Extrapolation of the fitted model (dotted red line) predicts that 41% of mammalian and 37% of primate CNEs would exhibit activity if sampled across an exhaustive range of tissues. **k)** Complexity and distribution of CNE activity patterns. Each gray dot represents an individual tissue-activity pattern. Pattern complexity is defined as the number of active tissues (*x*-axis; *n* = 1, 2, …, 19), with the *y*-axis showing the number of CNEs exhibiting each specific pattern for each complexity level. Red diamonds indicate the mean number of CNEs observed for each complexity level, while black circles represent the expected counts under a Binomial distribution. While the theoretical model exceeds the observed mean for low-complexity patterns (*n* = 1, 2, 3), the observed mean is higher for all levels where *n* ≥ 4. The characteristic U-shaped distribution underscores a functional dichotomy between highly tissue-specific CNEs and those exhibiting constitutive activity across all 19 tissues. See Supplementary Figure 4 for primate data. **ALT TEXT:** Graphs on functional characterization of CNEs, illustrating the distance of CNEs to TSS, their overlap with *cis*-regulatory elements identified by SCREEN, and the number of CNEs with DNase I hypersensitivity and H3K27ac epigenetic marks.

Regardless of their evolutionary age, all CNE sets exhibited significant enrichment for SCREEN-predicted candidate promoters and enhancers (*P*-values < 0.05, Fisher’s exact test; Supplementary Figure 1; Methods), highlighting their potential regulatory roles. In particular, of the mammalian CNEs, 34% overlapped with enhancers (Odds Ratio, OR = 3.8) and 3% with promoters (OR = 4.5). Primate CNEs showed slightly lower overlap rates (enhancers: 28%, OR = 3.3; promoters: 2%, OR = 3.3). Consistent with their genomic contexts, intronic CNEs were more frequently associated with promoters (OR = 6.2 for mammalian CNEs; OR = 4.1 for primate CNEs) than intergenic CNEs (OR = 1.7 for mammalian CNEs; OR = 1.4 for primate CNEs). Aligning with these overall findings, nearly all CNE sets demonstrated significant overlaps with SCREEN-predicted candidate promoters and enhancers across each of the 21 analyzed tissues and cell types (Methods), emphasizing the high regulatory potential of CNEs irrespective of the cellular context (Supplementary Figure 1).

While SCREEN predictions provide a useful categorical framework, they do not capture the full dynamic range of a sequence’s activity. To gain a more granular profile of chromatin accessibility and *cis*-regulatory activity across our CNEs, we quantified DNase I hypersensitivity and H3K27ac fold-enrichment across 19 of the same tissues and cell types (Methods). Initial analysis confirmed that both mammalian and primate CNEs are significantly enriched for these marks relative to random genomic regions (Figures 2b and c). Stratifying the CNEs into five categories based on signal intensity, ranging from “inactive” (Category 1) to “highly accessible”/”acetylated” (Category 5) (Supplementary Tables 2 and 3; Methods), enabled us to identify high-confidence *cis*-regulatory elements.

A substantial fraction of the CNEs exhibited an open chromatin configuration, with 42% of mammalian and 38% of primate CNEs reaching Category 4 or 5 in at least one tissue. Naturally, per-tissue accessibility was more restricted, ranging from 6% to 17% for mammalian and 5% to 14% for primate CNEs (Figure 2d). While approximately 15% of the CNEs were in an open chromatin configuration in strictly one tissue, a large fraction of CNEs appeared to have more ubiquitous roles, with the mammalian subset tending toward higher pleiotropy. Specifically, 20% of mammalian and 17% of primate CNEs were accessible in three or more tissues, and 1.5% of mammalian and 1% of primate CNEs demonstrated constitutive accessibility across all 19 tissues (Figure 2e). Consequently, mammalian CNEs were accessible in an average of 4.5 tissues compared to 4.1 of primate CNEs.

Also a considerable fraction of CNEs exhibited high or very high acetylation (Categories 4 or 5) in at least one tissue, though the lineage trend was reversed, with primate CNEs showing a slightly higher fraction (58%) than mammalian CNEs (56%). Per-tissue occupancy varied, ranging from 4% to 23% for mammalian and 4% to 21% for primate CNEs (Figure 2f). H3K27ac enrichment exhibited a more pronounced balance between tissue-specificity and ubiquity than was observed for accessibility. Remarkably, 22% of mammalian and 23% of primate CNEs displayed high or very high acetylation in strictly one tissue, while a similar proportion—23% and 24%, respectively—did so in three or more tissues (Figure 2g). Most notably, the ubiquitousness observed in chromatin accessibility did not extend to histone acetylation: irrespective of conservation level, only 0.1% of CNEs featured high or very high acetylation across all 19 tissues.

We observed a strong mutual dependency between chromatin accessibility and histone acetylation (Figure 2h and Supplementary Figure 2): Across all tissues, 60% of non-acetylated CNEs (Category 1) remained, on average, in a closed chromatin configuration, while CNEs in a closed chromatin configuration were almost never acetylated. Conversely, CNEs in an open configuration (Category 5) were four times (mammalian) or three times (primate) more likely to exhibit high acetylation (Category 5) than their closed counterparts, with a reciprocal trend observed for CNEs featuring high acetylation. This reciprocal enrichment suggests that for the vast majority of CNEs, accessibility and acetylation represent a single, coordinated regulatory state.

The co-occurrence of open chromatin and histone acetylation serves as a definitive hallmark of active *cis*-regulatory elements (Shlyueva et al. 2014). Based on this signature, 29% of mammalian and 26% of primate CNEs are active in at least one tissue (Supplementary Table 4), representing a pool of 172,264 CNEs, with 59% being primate-specific sequences. Across individual tissues, active fractions remained sparse but followed a clear differentiation gradient, ranging from 2% in pluripotent H1-hESCs to 12% in terminally differentiated myotubes, astrocytes, and neurons (Figure 2i and Supplementary Figure 3). Saturation analysis predicted that exhaustive tissue sampling would eventually reveal activity for 41% of mammalian and 37% of primate CNEs (Figure 2j; Methods). Interestingly, CNE activity patterns across the 19 tissues under investigation follow a remarkable, U-shaped distribution that deviates sharply from random expectation (Figure 2k and Supplementary Figure 4). Specifically, we observed high activity convergence at the extremes: thus, thousands of CNEs shared identical, specific activity patterns in 1 to 4 tissues, while a distinct CNE subset—262 mammalian and 440 primate CNEs—exhibited ubiquitous activity across all 19 tissues. In contrast, CNEs with activity patterns involving 5 to 15 tissues were mostly individual, suggesting that selection favors “extreme” regulatory strategies of either strict specificity or broad ubiquity.

Together, our findings indicate that nearly one 40% of CNEs potentially function as *cis*-regulatory elements, typically within restricted cellular contexts.

### CNEs are enriched within TAD centers and particularly concentrated in gene-poor TADs associated with developmental processes

Having established the enrichment of active *cis*-regulatory elements at CNEs, we next sought to determine the potential contribution of CNEs to the three-dimensional (3D) organization of the genome.

While CNEs demonstrated a significant, albeit modest, overlap with CTCF binding sites (0.5% to 2.5% across tissues; *P*-values < 0.05, Fisher’s exact test; Methods), their distribution within topologically associating domains (TADs) followed specific non-random patterns. Thus, the number of CNEs per TAD strongly correlated with TAD size, with average

Spearman’s ρ across tissues of 0.62 and 0.61 for mammalian and primate CNEs, respectively (Figure 3a and Supplementary Figure 5). This correlation was notably weaker when considering only active CNEs (ρ = 0.34 and 0.28, respectively). Furthermore, CNE densities decreased towards TAD boundaries (*P*-values < 0.003; Figure 3b and Supplementary Figure 6; Methods), a finding that is consistent with prior evidence that CNE cluster boundaries co-localize with TAD boundaries (Harmston et al. 2017). Interestingly, we found a strong inverse correlation between evolutionary constraint on noncoding sequences and gene content. Specifically, TADs exhibiting high densities of deeply conserved mammalian CNEs were significantly gene-poor compared to their low-density counterparts. Thus, depending on the tissue, low-density TADs averaged 7-8 genes, whereas high-density TADs contained only 3-4 genes (*P*-values < 0.0002, computed with one-sided Wilcoxon’s tests; Methods). Also gene densities were substantially lower in high-density TADs (0.51-0.65 genes per 100 kb) compared to their low-density counterparts (0.96-1.27 genes per 100 kb). For primate CNEs, the number of genes between high- and low-density TADs remained comparable (6-9 vs. 5-7 genes), but a trend emerged in their gene densities.

**Figure 3.**
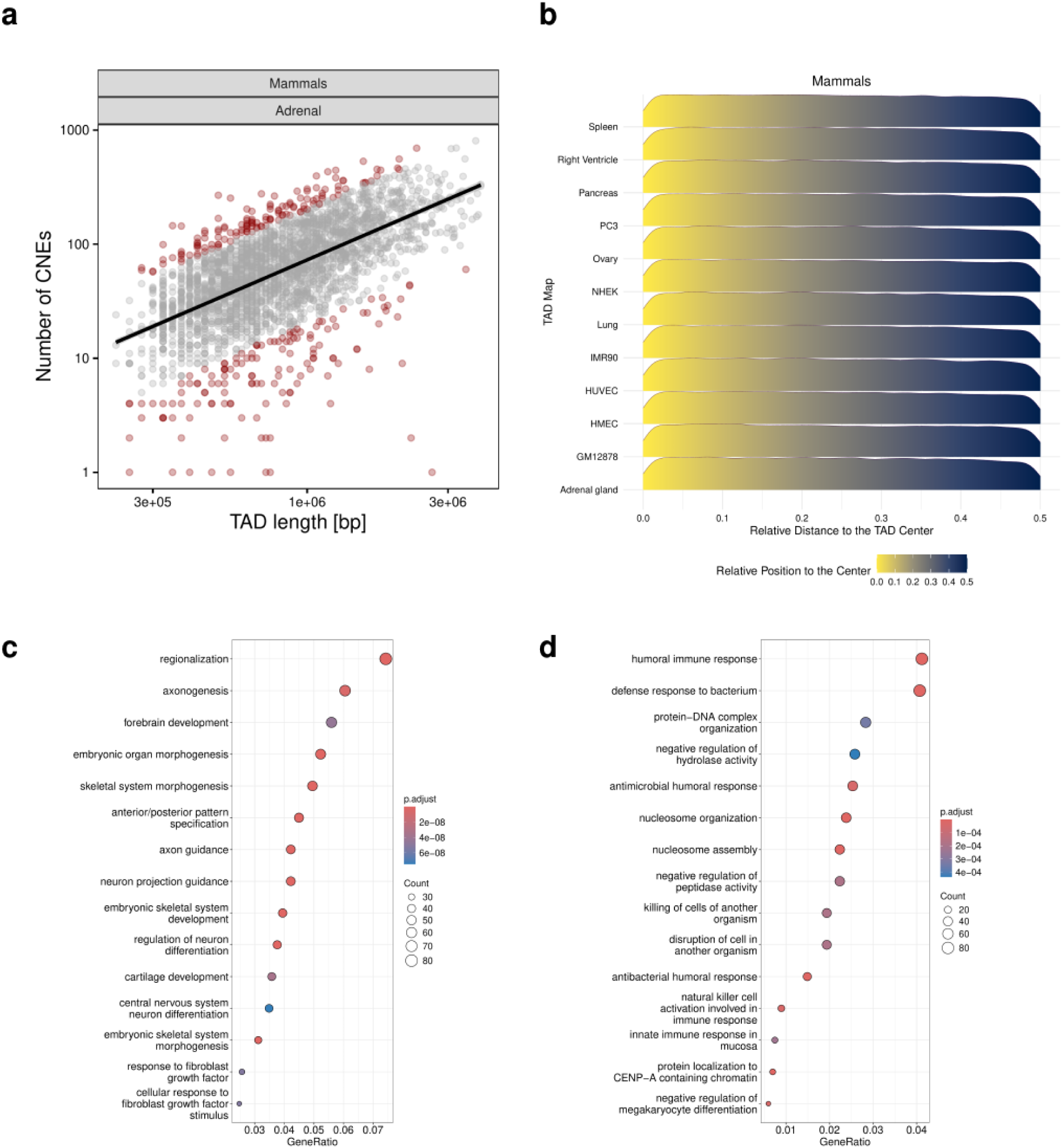
CNEs are enriched in TAD centers. **a)** Correlation between TAD length (in bp) and CNE density. Scatter plot illustrating the relationship between CNE number and TAD length, exemplified here for adrenal gland tissue and mammalian CNEs. The number of CNEs was strongly positively correlated with TAD length. TADs identified as outliers are highlighted in dark red. Specifically, outliers were defined as the 5% of TADs with the most extreme positive and negative residuals from a linear regression of CNE count against TAD length. See Supplementary Figure 5 for additional tissues and primate CNEs. **b)** Distribution of CNEs within TADs. Ridgeline plot showing the density distribution of mammalian CNEs relative to the center of their respective TADs. CNE density peaked at TAD centers and decreased significantly toward domain boundaries See Supplementary Figure 6 for primate data. **c**) and **d**) Functional enrichment analysis of CNE-dense and CNE-poor TADs, respectively. Gene Ontology (GO) terms for genes located within TADs partitioned by mammalian CNE density. TADs with high CNE density (c) are significantly enriched for genes involved in development and morphogenesis. Conversely, genes in TADs with low CNE density (d) are primarily associated with responses to external stimuli and environmental factors. Data for primate CNEs can be found in Supplementary Figure 7 and 8. Bubble size is proportional to the number of genes associated with each GO term, while the color gradient represents the statistical significance (adjusted *P*-value). **ALT TEXT:** Graphs on the intersection of CNEs and TADs, illustrating the distribution of CNEs within TADs and the gene ontology terms associated with genes in CNE-rich TADs compared to CNE-poor TADs.

Indeed, in TADs featuring high densities of primate CNEs, gene density ranged from 0.83 to 0.99 genes per 100 kb, compared to 1.07 to 1.26 in low-density TADs (*P*-values < 0.0002, computed with one-sided Wilcoxon’s tests; Methods). These observations corroborate previous studies showing that highly constrained regulatory regions preferentially co-localize with gene deserts (Ovcharenko et al. 2005). Furthermore, the genomic partition defined by CNE and gene densities also reflects functional modularization: genes in TADs with high CNE densities were primarily associated with developmental processes and morphogenesis (Figure 3c and Supplementary Figure 7); genes in TADs with low CNE densities were related to response to external factors like bacteria or chemokines or general immune system activation (Figure 3d and Supplementary Figure 8). Taken together, these results demonstrate that CNEs are organized within the 3D genome according to a distinct logic, which may either be required by their *cis*-regulatory roles or hint at other, potentially structural, roles.

### Network topology modulates the conservation of noncoding elements with *cis*-regulatory functions

Finally, we leveraged our dataset of active CNEs to systematically identify common gene misregulation mechanisms underlying complex human diseases. To assess the potential clinical relevance of CNEs, we first quantified their overlap with 364,201 GWAS-identified disease and trait variants (Methods). We found that the distribution of the variants across the CNEs in the dataset was highly skewed towards zero, with only 4% (7,527) of active CNEs harboring one or more variants. Out of the 955 phenotypes associated with at least 100 variants in the GWAS Catalog (Cerezo et al. 2025), we identified a significant overlap between active CNEs and the associated variants of 67% (636) of the phenotypes (Benjamini-Hochberg adjusted *P*-value < 0.05, computed with Fisher’s exact test), including schizophrenia (a disease), insomnia (a syndrome), and neuroticism (a personality trait) (Supplementary Figure 9). These neuropsychiatric phenotypes are known to involve substantial genetic overlap (Grotzinger et al. 2026) and common pathways related to neurodevelopment and synaptic signaling.

In total, 220 active CNEs harbored 223 SNPs associated with schizophrenia, insomnia or neuroticism. Overall, five of these elements harbored SNPs associated with two or more phenotypes (Figure 4a): one with schizophrenia and insomnia; another one with insomnia and neuroticism; and three with schizophrenia and neuroticism. For schizophrenia, about half (49%) of the 100 CNEs were mammalian-conserved and half (51%) were primate-conserved (Figure 4b). In contrast, the 67 CNEs linked to insomnia and the 58 CNEs related to neuroticism were mostly mammalian-conserved, at 66% and 59%, respectively. Putative promoters comprised 19 of the 220 CNEs, with 79% of them being mammalian-conserved. Specifically, eight serve as putative promoters for genes linked to schizophrenia (*C5orf63*, *FAM53C*, *FUT9*, *INHBE, SMG6, SPECC1*, *TAOK2, TCF20*), six for insomnia (*ADGRB3, NLGN1, ONECUT1, PHF21A, PPP1R18, SEMA6D*), and five for neuroticism (*ARPP21, CLIC1, CSNK1G1, RAI1,* Y*LPM1)*.

**Figure 4.**
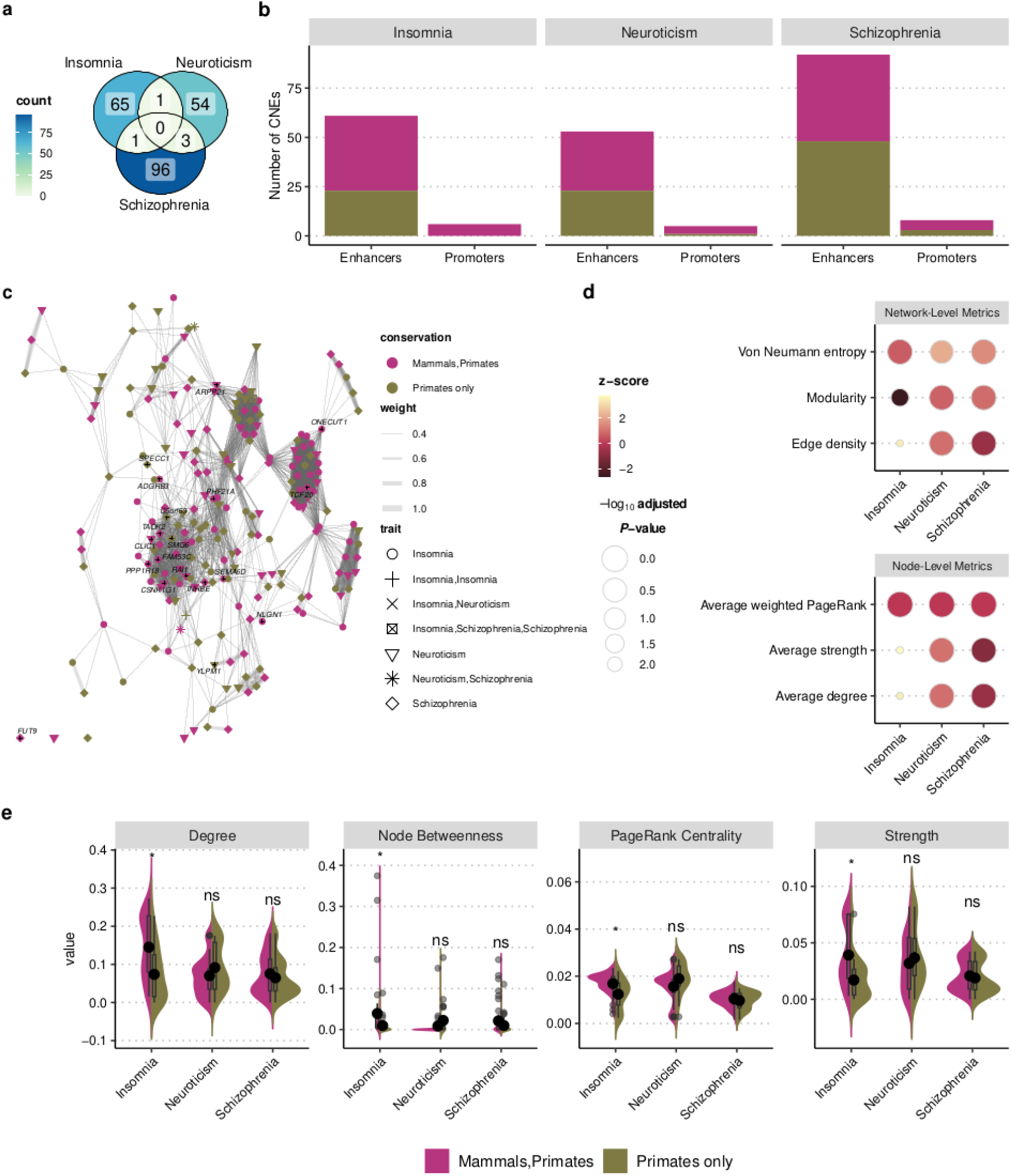
Network topology and evolutionary constraints of trait-associated CNEs. **a)** Venn diagram illustrating the overlap of CNEs harboring variants across the phenotypic traits schizophrenia, insomnia and neuroticism. Intersecting regions quantify the number of CNEs containing variants for multiple phenotypic traits. **b)** Distribution of CNEs annotated as promoters and enhancers harboring variants for each of the three analyzed phenotypic traits. Bar components differentiate the evolutionary depth of the CNEs, stratifying elements into those sharing deep mammalian-wide conservation (conserved in mammals and primates) versus those representing lineage-specific innovations (exclusively conserved in primates). **c)** Visual representation of the global network, comprising 220 nodes and 2,032 edges (isolated at an edge-weight threshold of 0.38). CNE nodes functionally annotated as promoters are marked with a small black “+” and labeled with their corresponding gene name. Edge thicknesses correspond to the pairwise tissue-activity Jaccard similarity index, visually distinguishing strongly from weakly synchronized CNE pairs. **d)** Dotplots displaying empirical *z-scores* for global network properties and aggregated node-level centrality metrics. *z-scores* represent the standard deviation shift of the observed network values relative to an empirical null distribution generated from 500 subnetworks randomly sampled from the global network matching the number of nodes in the subnetwork of interest and preserving the proportion of CNEs annotated as promoters and enhancers. Marker sizes are inversely proportional to Benjamini-Hochberg adjusted *P*-values derived from permutation testing, with asterisks indicating statistically significant deviations (adjusted *P*-value < 0.05) from random topological expectations. **e)** Centrality profile distribution of enhancer-annotated CNEs by evolutionary conservation. Violin plots displaying the topological property distributions, including degree, strength, weighted PageRank, and normalized betweenness centrality for CNEs annotated as enhancers. The split violin shapes contrast elements featuring deep mammalian-wide conservation against lineage-specific innovations (primate-specific). Embedded internal boxplots indicate the median and interquartile range for each distribution, while the overlaid dot-and-line markers represent the mean and standard deviation (± SD), illustrating the association between evolutionary constraint and topological importance of the CNEs. Statistical significance between conservation groups is indicated by asterisks (Benjamini-Hochberg adjusted *P*-value < 0.05; two-sided Wilcoxon rank-sum test). **ALT TEXT:** Graphs on the co-activation network inferred for active CNEs harboring variants associated with schizophrenia, insomnia, and neuroticism, illustrating different metrics of the global network and induced subnetwork, considering the conservation level of the CNEs.

To examine the evolutionary constraints on CNEs and as a case study, we constructed a coactivity network with nodes representing active CNEs harboring SNPs associated with schizophrenia, neuroticism, or insomnia and edges denoting co-activation (Methods). In this network, edge weights reflect the similarity of the activity profiles of the CNEs they connect. The original network comprised 220 nodes and 10,039 edges, forming a densely connected structure (Supplementary Table 5). Recursive weight pruning (Methods) showed that global connectivity was maintained by intermediate- and high-weight edges, while low-weight edges primarily contributed redundancy (Supplementary Figure 10). Thus, the network became increasingly modular but remained interconnected until ∼78% of edges were removed (weight threshold = 0.38); beyond this point the network fragmented, largely exceeding null expectations. The pruned network –which was used for all subsequent analyses– comprised 2,032 edges (Figure 4c). This network was sparsely connected (density = 0.08; average degree = 18.47) but retained high clustering (transitivity = 0.73) and a decentralized, uniformly distributed connectivity profile (von Neumann entropy = 5.08; Supplementary Table 6). Of the edges, 63% (1,283) linked CNEs harboring SNPs associated with different phenotypes, indicating substantial regulatory pleiotropy.

To investigate the regulatory architecture of phenotype-associated CNEs, we defined induced subnetworks for schizophrenia, insomnia, and neuroticism based on the CNEs harboring their respective variants (Methods). These subnetworks were benchmarked against size-matched subnetworks drawn from the network obtained for all three phenotypes, preserving the proportion of CNEs annotated as promoters and enhancers (Methods). Topological comparison revealed that the insomnia subnetwork represents an outlier compared to both the schizophrenia and neuroticism subnetworks (Figure 4d; Supplementary Table 7).

The insomnia subnetwork is characterized by a highly non-modular (average modularity = 0.55; *z-score* = −2.6) and hyper-connected (average degree = 8.03; *z-score* = +3.48) architecture, suggesting that it operates as a highly interdependent, single functional unit. Furthermore, this centralized network is dominated by highly overlapping tissue-activity profiles (average strength = 5.59; *z-score* = +3.79). As a result, the system is globally synchronized, with a large fraction of CNEs active within the same tissue contexts. Notably, CNEs exhibiting higher centrality metrics—degree, strength, weighted PageRank centrality and node betweenness—are associated with elements deeply conserved across mammals (Figure 4e). Indeed, because the system is globally synchronized, younger primate-specific innovations are relegated to the less critical periphery of the network.

The non-modular architecture of the insomnia subnetwork contrasts sharply with the modular topologies of schizophrenia and neuroticism. Both schizophrenia- and neuroticism-associated CNEs are partitioned into discrete, specialized functional teams (average modularities = 0.71 in both cases; *z-score* = +0.93 and *z-score* = +0.74, respectively). Furthermore, their elevated Von Neumann entropy values (4.23 and 3.62, respectively; *z-score* = +1.54 and *z-score* = +2.17) show that the CNEs in these subnetworks feature an overall wide breadth of activity across tissues. However, while the reduced average strength of schizophrenia (4.68; *z-score* = −1.03), reflecting low pairwise Jaccard similarity indices, highlights that this wide breadth is driven by individual CNEs active in single, but different tissues, the elevated average strength of neuroticism (3.45; *z-score* = +1.03) indicates that its breadth stems from CNEs that are simultaneously active across multiple tissues. The compartmentalization of these subnetworks into discrete modules relaxes the evolutionary constraints on individual sequences. Because mutations within an isolated module are unlikely to cause major disruptions across the broader network, purifying selection on these sequences is reduced, facilitating rewiring through sequence innovation. As a result, younger, primate-specific sequences are able to achieve high topological centrality within these systems.

Taken together, our integration of network biology and comparative genomics indicates that evolutionary constraint on *cis*-regulatory elements is not merely a reflection of its isolated, individual regulatory role. Instead, it reflects the topological context in which it is integrated, with specific structural demands and interconnectivity of the network dictating the level of evolutionary pressure exerted on the sequence.

## Discussion

CNEs have long been investigated for their involvement in gene regulation, most prominently as enhancers, yet their functional annotation across tissues and evolutionary scales remains incomplete. By leveraging large-scale epigenomic resources from ENCODE, we sought to provide a comprehensive characterization of CNE regulatory activity across diverse cellular contexts. Using phastCons elements of the 470-mammal alignment, we identified ∼700,000 autosomal CNEs, spanning ∼5% of the human genome. Of these, approximately ∼240,000 are conserved across all mammals, whereas ∼670,000 are conserved in the more recent primate lineage and encompass both ancestral sequences and lineage-specific gains. In both groups, ∼30% overlap SCREEN-predicted enhancers and ∼2% SCREEN-predicted promoters, supporting a predominant role as *cis*-regulatory elements. Providing further evidence for this, analysis of ENCODE DNase-seq and H3K27ac ChIP-seq data across 19 tissues revealed that ∼28% of CNEs are active as putative enhancers or promoters –defined by concurrent chromatin accessibility and H3K27 acetylation in at least one tissue–, while ∼0.1% are constitutively active across all tissues. Furthermore, modeling indicated that ∼50% would remain inactive across additional cell types, effectively characterizing a large fraction of CNEs as enhancers. This regulatory potential was distributed non-randomly across TAD architecture: CNEs clustered within TAD centers. Moreover, high CNE densities corresponded to gene-poor TADs comprising genes specialized in developmental processes. Finally, our co-activation network analysis suggests that evolutionary conservation reflects systemic constraints on the regulatory network’s architecture. Thus, purifying selection would preserve sequences based on their indispensable topological role rather than their intrinsic properties.

While our analysis framework provides a robust basis for assessing the link between sequence conservation and *cis*-regulatory function, several inherent constraints warrant careful consideration. First, our CNEs are based on phastCons elements derived from a multiple alignment between 470 mammalian assembly sequences. Although we define “mammalian” and “primate” CNEs as distinct sets, they represent evolutionary nested groups, with the mammalian CNE set being largely recapitulated within the primate CNE set. Notably, primate CNEs are not necessarily strictly restricted to the primate lineage; rather, they identify regions where the alignment provides sufficient signal to confirm conservation within primates. Furthermore, the varying quality of the assemblies, particularly those representing initial species drafts, may lead to an underestimation of the number of CNEs in each lineage. To reduce the impact of low-quality data, we only included assemblies with an N50 > 100 kb in the ancestral reconstruction. In addition, the uneven sampling bias across the phylogeny may have introduced a bias towards specific clades. For example, while 94 Glires assemblies account for only 4% of the group, seven assemblies represent 35% of Xenarthra. Equally important, because the underlying alignment is human-centric, our findings are inherently biased toward elements maintained in the human lineage. Moreover, the functional characterization of our CNEs relied on ENCODE. We recognize that the integration of alternative data repositories or independent primary studies would likely offer additional insights into the cell-type-specific activities of these elements. However, we purposely opted to utilize this unified, high-quality resource to ensure consistent standards and circumvent the confounding influence of batch effects inherent in separate primary sources (Leek et al. 2010). Beyond data selection, our definition of *cis*-regulatory activity relied on a specific enrichment threshold (*log*_2_ *FC* = 1. 5). Finally, while our functional characterization of CNEs rests on a solid framework, prospective validation through functional assays will be required to definitively confirm the accuracy of the inferences.

Our results underscore a striking symmetry: While approximately half of ENCODE’s candidate *cis*-regulatory elements are conserved across mammals (Andrews et al. 2023), our saturation analysis suggests a comparable proportion of CNEs function as active *cis*-regulatory elements across a comprehensive range of cellular contexts. This is consistent with the fact that 45% of highly conserved human-fish sequences have been validated as functional enhancers in transgenic mouse assays (e.g., (Pennacchio et al. 2006)). In contrast, an earlier extrapolation based on DNase-I hypersensitivity, histone modifications, and transcription factor binding peaks across 10 human ENCODE tissues and cell types [PMID: 22684627] projected that ∼85% of CNEs would exhibit *cis*-regulatory activity. This discrepancy can be attributed to a fundamental difference in methodology and biological definition. Crucially, those findings rely on single peak overlaps—sometimes as minimal as 1 bp—a relatively lenient metric. Indeed, if we defined activity simply as the presence of either DNase-I hypersensitivity or H3K27ac signal, our estimates would also increase substantially, up to 63-65%. Furthermore, such peaks often represent opportunistic chromatin accessibility or transient TF occupancy rather than established *cis*-regulatory activity (e.g., (Bozek and Gompel 2020)). Instead, we defined *cis*-regulatory activity based on the co-occurrence of DNase hypersensitivity I and H3K27ac, for which we require a minimum *log*_2_ 1.5 fold-enrichment. Naturally, the choice of threshold naturally influences these proportions; for instance, relaxing the requirement to *log*_2_ 1.25 increases the active fraction to 34% (mammalian) and 30% (primate), while a more stringent *log*_2_ 1.75 reduces them to 26% and 22%, respectively. Still, the co-occurrence of substantial levels of DNase-I hypersensitivity or H3K27ac is widely accepted as a high-confidence indicator of regulatory function (e.g.,(Zhang et al. 2013)), and should therefore provide a more conservative, biologically grounded estimate. Also, it is important to note that since our saturation analysis is founded on tissues and cell lines as diverse as the adrenal gland and heart, it is inherently ill-equipped to capture highly specific spatiotemporal *cis*-regulatory activity. This reflects a historical hurdle in the field: in 2007, a landmark study deleted four noncoding UCEs in mice, concluding to the community’s surprise that the animals remained phenotypically normal (Ahituv et al. 2007). It took over a decade to resolve this apparent paradox—the existence of sequences under extreme purifying selection that seemingly lacked function—by eventually revealing that the absence of these elements causes subtle but critical defects in specific neuronal populations (Dickel et al. 2018). Because such CNEs act as “fine-tuners” that operate within niche spatiotemporal windows, they would be expected to lack substantial DNase-I hypersensitivity or H3K27ac signatures in the majority of tissues, remaining invisible to bulk tissue saturation analysis. Consequently, our results provide a robust estimate of constitutive or broad-acting *cis*-regulatory CNEs, effectively reconciling biochemical saturation with known evolutionary and experimental constraints.

Beyond acting as enhancers and promoters, CNEs could fulfill other *cis*-regulatory roles, for example as insulators —estimated to account for roughly 2% of human CNEs (Xie et al. 2007)— or as transcriptional repressors. While individual histone modifications, such as H3K27me3 facultative heterochromatin and H3K9me3 for constitutive heterochromatin (Wu et al. 2023), have been described to delineate these functional classes, their global cataloging remains relatively limited, and the precise architectures of active insulators and repressors are not yet fully understood. In addition, CNEs likely serve a diverse array of functions. Supporting this possibility, previous estimates indicate that 11% of human-mouse CNEs overlap with predicted Scaffold/Matrix Attachment Regions (S/MARs), suggesting a role in facilitating higher-order chromosomal architecture (Glazko et al. 2003). Furthermore, CNE clusters have been spatially associated with TADs (Harmston et al. 2017) and may constitute tethering points for TAD boundaries (Polychronopoulos et al. 2017). Alternatively, a substantial fraction of CNEs may be unannotated noncoding RNAs (ncRNAs). Indeed, approximately 20% of mammalian CNEs are presumed to represent unannotated ncRNAs (Hemberg et al. 2012). Given that the vast majority of human long non-coding RNAs (lncRNAs) likely remain uncharacterized (Iyer et al. 2015), it is plausible that many CNEs correspond to lncRNAs or lesser understood ncRNA classes. Finally, some CNEs may reflect functional decay and represent ancestral *cis*-regulatory elements that have undergone functional attenuation. This hypothesis is supported by evidence that ∼32% of CNEs exhibit significant substitution rate variations across lineages (Kim and Pritchard 2007), indicating that a substantial portion of CNEs are undergoing functional decay, likely driven by regulatory rewiring or redundancy. These possibilities, however, are by no means exhaustive.

Although only a modest fraction of GWAS variants mapped to CNEs, these overlaps were significantly enriched across thousands of phenotypes, with particularly strong signals for three neuropsychiatric phenotypes: schizophrenia, insomnia, and neuroticism. The phenotypes are known to share genetic aetiology and overlap in neurodevelopmental and synaptic pathways, while representing distinct levels of phenotypic affection: a disease, a syndrome, and a personality trait, respectively. CNEs harboring variants associated with these phenotypes are enriched for deep mammalian conservation. While primate-specific sequences constitute the majority (59%) of all active CNEs across the 19 tissues under investigation, this trend is inverted among risk loci: 56% of phenotype-associated CNEs are conserved in mammals, with only 44% being primate-specific. This enrichment for ancestral sequences is particularly stark among promoters, where an extreme majority—15 out of 19 elements—are mammalian-conserved.

To understand the relationship between evolutionary constraint and regulatory context, we constructed a co-activation network comprising the 220 active CNEs that harbor SNPs associated with any of these three phenotypes. Rather than functional redundancy, co-activity represents spatial synchronization, as these CNEs are likely to have different gene targets. Because many of these CNEs are active across multiple tissues –they are pleiotropic– they are under an additive layer of purifying selection because a mutation would cause widespread, multi-system disruptions, explaining why they are enriched for deep mammalian conservation in the first place. Furthermore, this enrichment is consistent with the evolutionary history of the gene regulatory networks underlying these phenotypes. For instance, the genetic core of insomnia maps to master circadian pacemakers (CLOCK, PER, CRY) and fundamental neurotransmitter systems–such as such as gamma-aminobutyric acid (GABA) and melatonin–that remain highly conserved from invertebrates (e.g., *Drosophila*) through humans (Brown et al. 2019). Also the network underlying neuroticism is anchored to an ancient threat-detection mechanism that enables mammals to rapidly perceive danger and trigger immediate physiological survival responses —relying on conserved networks involving the amygdala, hippocampus, and prefrontal cortex (Tzschoppe et al. 2014). And even the schizophrenia gene regulatory network—despite its highly complex human manifestation—shares fundamental mammalian components, such as basic N-methyl-D-aspartate (NMDA) and GABAergic synaptic transmission (Gaspar et al. 2009). Crucially, the varying degrees of recent primate expansion are reflected in the fractions of deeply conserved mammalian elements across these phenotypes. Thus, while 66% of CNEs harboring insomnia-associated SNPs are conserved across mammals, this fraction decreases to 59% in neuroticism and to 49% in schizophrenia. Indeed, as primates developed complex social hierarchies, the regulatory networks controlling cortical inhibition evolved rapidly to accommodate higher-order traits—such as social anxiety and environmental appraisal—that define the neuroticism spectrum (Latzman et al. 2018). Likewise, schizophrenia is widely understood as a byproduct of a massive, rapid brain expansion within primate lineages that relied on rewiring the original mammalian network (Hanson et al. 2025).

Our results demonstrate that network topology shapes the evolutionary pressure experienced by *cis*-regulatory elements. In highly centralized, non-modular networks like insomnia, structural burden—the degree to which downstream network stability and functional integrity depend on a single node—is heavily concentrated within a tightly interconnected core; because a mutation within an element in this core has the potential to destabilize the entire system, it is subjected to intense purifying selection. Consequently, highly central elements in the insomnia subnetwork are predominantly mammalian-conserved. Conversely, decentralized, modular networks distribute this structural burden across functionally discrete modules, thereby accommodating evolutionary innovation. This structural divergence accounts for the subtle differences in centrality values observed between mammalian and primate-specific elements within the schizophrenia and neuroticism subnetworks. Interestingly, despite their shared modularity, these subnetworks exhibit opposing internal trends that highlight how a sequence’s conservation is dictated by its precise regulatory context rather than its isolated function. Compared to schizophrenia, primate-specific CNEs in the neuroticism subnetwork are associated with higher strengths.

This divergence directly reflects the distinct neurobiological architecture of these two phenotypes: while neuroticism is a complex behavioral trait localized within neocortical regions responsible for high-order cognitive processing (Carleton 2016), schizophrenia is a systemic neurodevelopmental disorder characterized by disrupted synaptic signaling and diffuse, global brain dysconnectivity (Pearlson 2000). Because neuroticism is driven by recent neocortical expansion, the primate-specific elements themselves constitute the regulatory architecture that overlays the ancestral mammalian neocortex; the regulatory burden shifts to these young elements as they mediate these newly acquired cognitive functions. In contrast, schizophrenia targets global brain development and synaptic plasticity—foundational processes optimized early in mammalian evolution and maintained under intense negative selection. Because central disruptions to the underlying pathways are highly deleterious, primate-specific innovations are restricted to the periphery. Ultimately, these observations underscore that evolutionary conservation is not merely an intrinsic property of a *cis*-regulatory element’s isolated function, but a consequence of its topological neighborhood and role within the network in which it is embedded.

In summary, we present a comprehensive catalog of CNEs that integrates evolutionary constraint with biochemical evidence for regulatory activity, providing a robust framework for investigating the conserved core of the human *cis*-regulatory landscape. This resource facilitates the systematic study of a regulatory logic that has remained remarkably stable through the diversification of mammalian and primate lineages. By reconciling genomic conservation with active molecular signatures across a diverse tissue panel, we estimate that approximately 40% of CNEs function as *cis*-regulatory elements, with activity predominantly restricted to a discrete number of tissues. In addition, our findings highlight an association between evolutionary conservation and higher-order chromatin organization. Finally, we present evidence that regulatory network topology and an element’s structural burden within its specific network fundamentally modulate its evolutionary conservation, highlighting how this context is essential to understanding observed patterns of constraint.Decades after the completion of the Human Genome Project, the functional annotation of the noncoding genome remains a challenge, reinforcing the need for more comprehensive and granular functional resources. Ultimately, characterizing CNEs is an important step toward understanding the interplay between functional integrity and regulatory innovation in the human genome. This is fundamental for elucidating the molecular basis of human health and disease.

## Materials and Methods

### Conserved Element Identification

PhastCons elements derived from a multiple sequence alignment of 470 mammalian genomes to the human reference genome assembly (GRCh38/hg38) were downloaded from the UCSC Genome Browser (https://hgdownload.cse.ucsc.edu/goldenPath/hg38/phastCons470way/hg38.phastConsElem <u>ents470way.bb</u>, last accessed on June 11, 2026; (Perez et al. 2025)). The bigBed file was converted to bed format using UCSC Kent tools bigBedToBed (v357, (Kuhn et al. 2013)). It contained more than 30 million phastCons elements, of which 29 million were located on chromosomes 1-22, X or Y and retained for further analyses. PhastCons elements ranged in length between 1 bp and 7,144 bp, with a median of 4 bp.

### Conserved Element Annotation

The GENCODE v45 (Mudge et al. 2025) annotation of the human reference genome (GRCh38/hg38) was downloaded from https://ftp.ebi.ac.uk/pub/databases/gencode/Gencode_human/release_45/gencode.v45.chr.a <u>nnotation.gff3.gz</u> (last accessed on March 25, 2024).

Conserved elements were classified as “coding” if at least 30% of their length overlapped protein-coding sequences (represented as “CDS” features in the annotation file), or if they encompassed a complete protein-coding sequence. The remaining elements were defined as conserved noncoding elements (CNEs). Genomic overlaps were identified using BEDTools intersect (v2.30.0, (Quinlan and Hall 2010)) with the “-wao” option.

CNEs exhibiting any degree of overlap with the coordinates defining “gene” features within the reference genome annotation were classified as “intronic”; the remaining CNEs were considered “intergenic”.

### Purifying Selection of CNEs in Humans

Common human SNPs were downloaded from the UCSC genome browser (https://hgdownload.soe.ucsc.edu/goldenPath/hg38/database/snp151Common.txt.gz).

BedTools intersect was used to count the number of SNPs in CNEs and random genomic regions.

### Generation of Random Genomic Regions Corresponding to CNEs

The genomic coordinates of the CNEs were used to generate a size- and count-matched set of random genomic loci with BEDTools shuffle for comparative analysis. Shuffling was restricted to the chromosomes containing the features of interest (“-chrom”).

### Evolutionary Conservation of Conserved Elements

Average phastCons scores were computed based on https://hgdownload.cse.ucsc.edu/goldenPath/hg38/phastCons470way/hg38.phastCons470w ay.bw (last accessed on March 4, 2026) using the pyBigWig python package (v0.3.23, https://github.com/deeptools/pyBigWig, last accessed on March 4, 2026).

### Identification of CNE Orthologs

Multiple sequence alignments were obtained from https://hgdownload.soe.ucsc.edu/goldenPath/hg38/multiz470way/maf/ (last accessed on March 4, 2026). Only assemblies (Supplementary Table 8) with an N50>100 were used for the analysis. In cases where a species was represented by multiple assemblies within the alignment, the assembly exhibiting the highest N50 value was chosen. This procedure resulted in 201 assemblies. Chromosome/scaffold information (name and size) for the assemblies was either retrieved directly from the “chromInfo” tables hosted by the UCSC genome browser (e.g. http://hgdownload.soe.ucsc.edu/goldenPath/hg38/database/chromInfo.txt.gz; last accessed on March 4, 2026) or with the according GCA-ID from NCBI database (e.g. https://api.ncbi.nlm.nih.gov/datasets/v2alpha/genome/accession/GCA_006542625.1/downlo <u>ad?include_annotation_type=SEQUENCE_REPORT</u>; last accessed on March 4, 2026). Or for assemblies, where the fasta file had to be downloaded from the DNA ZOO project (e.g. https://dnazoo.s3.wasabisys.com/Macaca_fuscata/Macaca_fuscata_HiC.fasta.gz; last accessed on March 4, 2026), the chromosome/scaffold length file was generated using faidx (v1.19.2) from samtools (Danecek et al. 2021).

Alignment blocks containing the CNEs were extracted and processed using MafFilter (v1.3.1, https://jydu.github.io/maffilter/, last accessed on March 4, 2026) with parameter “input.dots=as_gaps”. Specifically, the ExtractFeature module was used to filter and trim alignment blocks, retaining only those overlapping with the CNEs. CNEs were defined as features in a GTF file. The ExtractFeature module was configured with “ref_species=hg38”, “feature.format=GTF”, “feature.file.compression=none”, feature.type=all”, “complete=no” (allowing partial overlaps), and “ignore_strand=no” (enforcing strand-specific overlaps). Subsequently, the RemoveEmptySequences module was applied to remove any sequences within blocks consisting solely of gap characters (“unresolved_as_gaps=no”). Resulting alignment blocks were written with the OutputAlignments module to a compressed file, unmasked (“compression=gzip, mask=no”). Extracted MAF blocks were reformatted into a custom BED-like representation. Each record began with the human reference sequence chromosome, start, end, and strand. These fields were followed by comma-separated fields providing details of the alignment block: the assembly identifiers of all aligned sequences, their corresponding chromosomes, their start coordinates, their aligned sequences, and their strands. Negative strand coordinates were reverse-complemented according to the chromosome sizes of the corresponding assemblies. BEDTools intersect was then used to identify the MAF blocks whose reference sequence coordinates were fully or partially contained within each CNE. Next, for each CNE and each non-reference sequence in a MAF block associated with that CNE, we merged alignment segments separated by at most 5 bp in the corresponding non-reference sequence (i.e., due to insertions in the reference sequence or in another non-reference sequence in the MAF, or to local inversions in the non-reference sequence of interest) to represent single, continuous segments. Finally, the coordinates of the non-reference sequence corresponding to the CNE were obtained by clipping the reference sequence in the MAF block to the precise boundaries of the CNE. For stringency, we retained only the ortholog with the greatest number of bases in the alignment relative to the base of the CNE; further, we required the number of bases in the alignment of the corresponding ortholog to exceed 24 bp.

### CNE Ancestral Reconstruction

The CNEs and their orthologs together with the phylogenetic tree corresponding to the Hiller Lab 470 Mammals alignments (https://hgdownload.soe.ucsc.edu/goldenPath/hg38/multiz470way/hg38.470way.nh, last accessed on June 11, 2026) was used to infer the ancestral presence/absence of each CNEs using the ace() function of the “ape” R package (v 5.8-1, (Paradis and Schliep 2019)) with a discrete model (option type=”discrete”).

### Overlap with Transcription Start Sites and Functional Enrichment Analysis of Genes Nearest to CNEs

Transcription start sites (TSS) for hg38 were downloaded from the Ensembl Biomart database (Dyer et al. 2025). BEDTools intersect was employed to identify CNEs enclosing a TSS. BEDTools closest was used to identify the nearest TSS to each CNE. The gene ontology (GO) terms associated with the genes corresponding to those TSS were compared between mammalian and primate CNEs with the “clusterProfiler” R package (v4.14.6, (Yu et al. 2012)).

### Overlap with Candidate Regulatory Elements Predicted by SCREEN

Enhancers and promoters predicted by SCREEN V3 (ENCODE Project Consortium et al. 2020) were obtained from https://screen.encodeproject.org (last accessed on June 11, 2026; Supplementary Tables 9 and 10). Overlaps with CNEs were assessed with BEDTools intersect and BEDTools fisher.

### Identification of Functionally Active CNEs

DNase-seq and H3K27ac ChIP-seq BAM files, and for ChIP-seq corresponding control BAM files, were obtained from the ENCODE (Luo et al. 2020; Kagda et al. 2025) data repository for the same tissues for which SCREEN predictions were available (Supplementary Tables 11 and 12). DNase-seq BAM files were converted to signal density tracks (bigWig) using deeptool’s bamCoverage with parameters “--binSize 10 --extendReads 300 --normalizeUsing RPKM”. H3K27ac ChIP-seq BAM files were processed alongside their respective controls with deeptool’s bamCompare with parameters “--binSize 10 --extendReads 300 --scaleFactorsMethod SES --skipNonCoveredRegions”. Average DNase and H3K27ac signals across each CNE were calculated from the bigWig files with the pyBigWig Python module (v0.3.23, https://pypi.org/project/pyBigWig/, last accessed on June 11, 2026).

For each tissue, CNEs were assigned to one of five categories based on their DNase-seq *log*_2_ enrichment over background and their H3K27ac *log*_2_ enrichment was calculated as the ratio of a CNE’s average DNase-seq signal to the mean signal of size-matched random regions. The following thresholds were applied to both metrics: Category 1: value < 0.5; Category 2: 0.5 *≤* value < 1; Category 3: 1 *≤* value < 1.5; Category 4: 1.5 *≤* value < 2; and Category 5: value > 2. CNEs assigned to Categories 4 or 5 based on their DNase-seq *log*_2_ enrichments and Categories 4 or 5 according to their H3K27ac *log*_2_ FCs were considered functionally “active”.

### DNase I hypersensitivity and H3K27ac Enrichment Analysis

Average DNase I hypersensitivity and H3K27ac signals for mammalian and primate CNEs were compared against size-matched random genomic regions. For each CNE/random genomic region, the maximum signal across all surveyed tissues was identified. We then calculated the observed proportion of CNEs (*p_f_*) and random regions (*p_b_*) exceeding a sliding signal threshold across the entire dynamic range. Enrichment was defined as the fold change between these two proportions 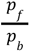. To ensure the reliability of the enrichment estimates, we filtered for results where the standard error (SE) of the *log*_2_ FC was less than 0.15. This SE was calculated as 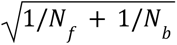, where *N*_*f*_ and *N*_*b*_ are the numbers of CNEs and random regions above the threshold, respectively.

### CNE Activity Saturation analysis

To estimate the total pool of active CNEs across an expanding number of tissues, we employed a permutation-based approach followed by non-linear regression. The columns (tissues) of the original binary matrix representing CNE activity were shuffled 100 times to generate a null distribution, and for each iteration, we calculated the cumulative number of active CNEs observed after sequentially including *n* tissues, where *n* = 1, 2, …, 19. For each tissue increment *n*, we computed the mean number of active CNEs across all 100 iterations.

To provide robust starting values for the maximum capacity (*V_max_*) and the Michaelis constant (*K_m_*) for the non-linear estimation, we performed a Lineweaver-Burk transformation using the lm() function in R on the reciprocal of the data. Finally, we fitted a Michaelis-Menten saturation model to the mean number of active CNEs using the nls() function in R, following the formula:

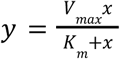

where *y* represents the cumulative number of active CNEs and *x* represents the number of sampled tissues.

### CNE Overlap with TADs

TAD coordinates were downloaded from the 3D Genome Browser https://3dgenome.fsm.northwestern.edu/; last accessed on March 4, 2026) for right ventricle, spleen, pancreas, ovary, NHEK, Lung, PC3, IMR90, HUVEC, HMEC, GM12878, and adrenal gland (Supplementary Table 13). The interquartile range (IQR) method was applied to discard TADs with outlier sizes from each dataset before proceeding with the analysis. Overlaps between CNE and TAD coordinates were assessed using BEDTools intersect.

### CNE Distribution within the TADs

Then we evaluated if the center 30,000 nucleotides region of the TADs contained statistically more CNEs than 30,000 nucleotides (half inside the TAD, half outside) at the boundary.

### Identification of TADs with Low and High CNE Density

A linear regression was performed to assess the dependency of the number of CNEs per TAD on TAD size. TADs with unusually high or low number of CNEs were defined as those with the largest deviations from the fitted line, specifically selecting the TADs corresponding to the 5% most positive and 5% most negative residuals.

### Functional Enrichment Analysis of Genes in TADs with Low and High CNE Density

TSS of protein-coding genes in TADs with low and high CNE density were identified with BEDTools intersect. Functional enrichment analysis of the corresponding genes was carried out with the clusterProfiler R-package (v4.14.6, (Yu et al. 2012)).

### Identification of CTCF-associated TAD Boundaries

CTCF sites were downloaded as BED files from the ENCODE data repository. TAD boundaries were defined as the genomic region corresponding to the meeting ends of two adjacent TADs after symmetrically extending it by ± 25,000 bp from its start and end coordinates. BEDTools intersect was used to identify TAD boundaries including CTCF sites.

### Retrieval of Disease/Trait-associated Variants

Disease and complex-trait associated variants were retrieved from the GWAS Catalog (v1.0; https://www.ebi.ac.uk/gwas/api/search/downloads/full; last accessed on March 4, 2026).

Variants associated with schizophrenia, insomnia, and neuroticism were retrieved from the GWAS query interface using Experimental Factor Ontology (EFO) identifiers: MONDO_0005090 (schizophrenia), EFO_0004698 (insomnia), and EFO_0004257 and EFO_0007660 (neurotic disorder and neuroticism measurement, respectively). This approach ensures a coherent consideration of synonyms. Child trait data were included.

### CNE-based Disease Regulatory Network Construction

Activity profiles of CNEs harboring GWAS variants associated with schizophrenia, neuroticism, and insomnia were compiled across 19 tissues. For each tissue, only CNEs reaching category 4 or 5 for DNase I hypersensitivity fold-enrichment and H3K27ac fold-enrichment in at least 5 tissues were classified as “active”. Pairwise coactivation was quantified using the Jaccard similarity index, calculated as the complement of the Jaccard dissimilarity returned by the vegdist(method=”jaccard”) function of the vegan R package (v2.7-2; (Oksanen et al. 2026) with “method=”jaccard””. The Jaccard metric was selected over correlation-based approaches because it captures shared activation while excluding co-inactivity, which dominates zero-inflated chromatin accessibility datasets.

A single network incorporating CNEs from all three phenotypes was constructed using the igraph R package (v2.2.1; (Csárdi and Nepusz 2006)). Specifically, the network was generated from the distance matrix via the graph_from_adjacency_matrix() function, enforcing an undirected structure (mode = “undirected”), assigning Jaccard similarity indices as edge weights (weighted = TRUE), and omitting self-loops (diag = FALSE).

### Network Robustness Analysis and Filtering

To evaluate the effect of low-weight edges in network architecture, we performed a percolation analysis in which edges were iteratively removed in order of increasing weight across 100 pruning levels, corresponding to increasing proportion of the total number of edges. At each pruning step, we quantified the number of connected components, the number of isolated nodes (degree=0), the relative size of the largest connected component, and louvain modularity of the largest connected component. Outcomes were benchmarked against a null model in which the same proportion of edges was randomly removed at each pruning level. Network metrics were summarized across 100 independent iterations. The results of the percolation analyses were used to establish a weight threshold to prune uninformative edges from the network.

### Network Description

Network topology was characterized using functions from the igraph R package.

Local connectivity was assessed based on node degree (the total number of edges connected to a node) and strength (the sum of all edge weights connected to a node) with the degree() and strength() functions, respectively. Node importance was captured using weighted PageRank centrality, via the page_rank() function. The central role of nodes in bridging different parts of the network was measured using normalized weighted betweenness centrality with the betweenness() function. For this calculation, the graph was treated as undirected (”directed = FALSE”) and scores were scaled between 0 and 1 (”normalized = TRUE”). Furthermore, because the betweenness() function interprets weights as distance rather than similarity, the inverted edge weights were passed to the “weights” argument so that stronger connections were treated as the shortest paths.

Network modules were identified using the Louvain optimization algorithm implemented in the cluster_louvain() function. Modularity values were extracted with the modularity() function. Overall connectivity was assessed by calculating network density (edge_density()) and transitivity (transitivity()).

Differences in node-level metric distributions were assessed using Kruskal-Wallis rank sum tests followed by post-hoc pairwise Wilcoxon rank-sum tests with Benjamini–Hochberg correction, as implemented in the stats R package (R v.4.4.1). The spectral complexity of the network was quantified using Von Neumann entropy derived from the eigenvalues of the graph Laplacian, obtained using igraph’s laplacian_matrix() function and the eigen() function from the base R package.

### Comparative Analyses of Phenotype-specific Subnetworks

Induced subnetworks were generated by retaining only phenotype-specific nodes from the full network.Metrics for induced subnetwork were compared to a null distribution of 500 subnetworks randomly sampled from the global network matching the number of nodes in the subnetwork of interest and preserving the proportion of CNEs annotated as promoters and enhancers. Empirical *P*-values were determined by comparing observed metrics against the null distributions. Effect sizes were summarized as *z*-scores, defined as the number of standard deviations the observed value shifted from the mean of the null distribution.

When comparing the network topology of mammalian and primate-specific CNEs, we focused solely on CNEs annotated as enhancers. CNEs annotated as promoters were excluded due to their stronger conservation and their characteristically high network strength, which would distort connectivity metrics.

## Data availability statement

The data and sources underlying this article are provided within the manuscript or supplementary information files.

