## Supplementary material for "Multi-layered characterization of ∼700,000 conserved noncoding elements within the human genome"

### Table of Contents

**Supplementary Figures and Legends** **2**

**Supplementary Tables and Legends** **14**

### Supplementary Figures and Legends

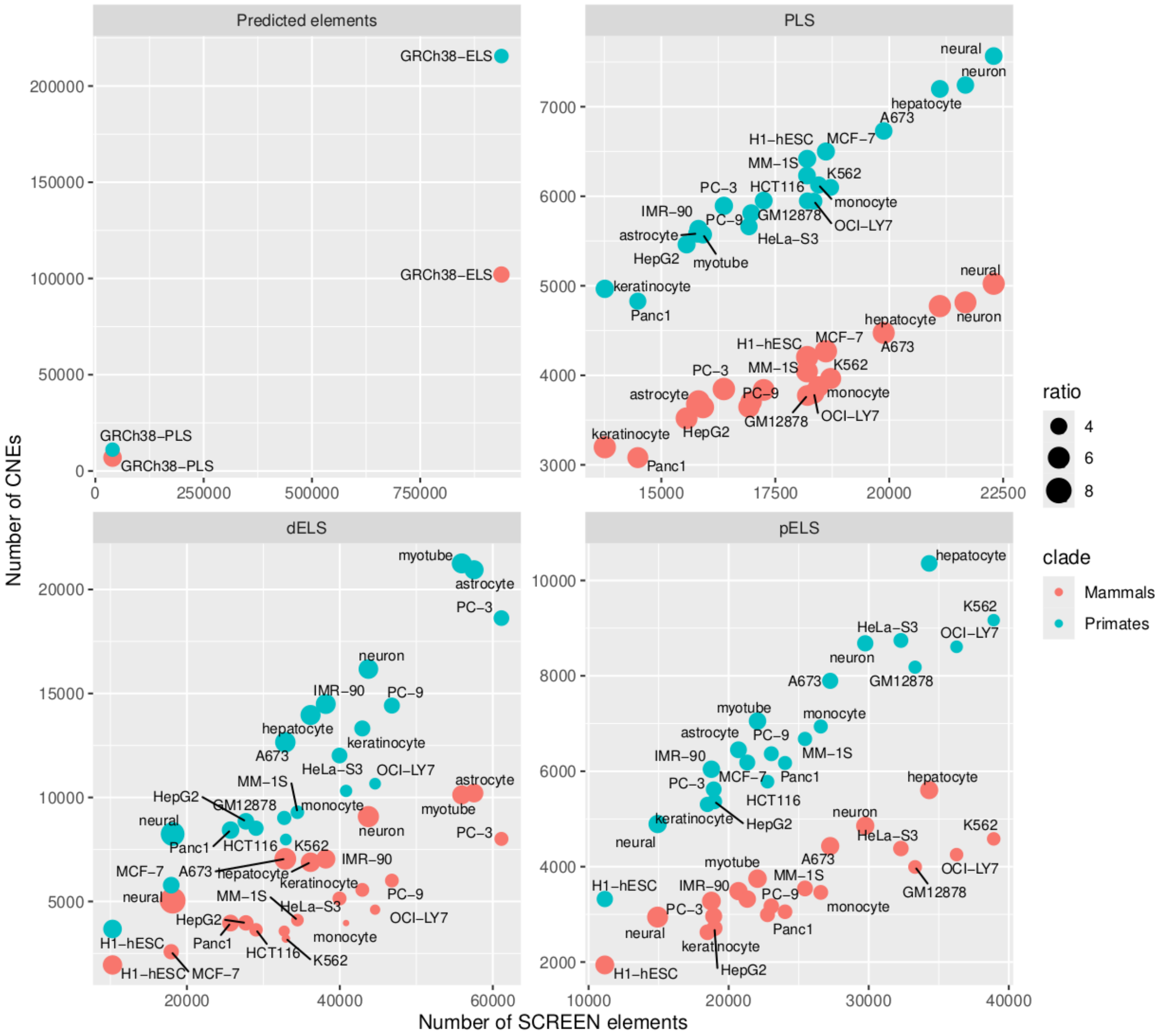


**Supplementary Figure 1. Overlap of CNEs with functional elements predicted by the SCREEN database.** Scatter plot with the number of overlapped CNEs on the y-axis and the number of overlapped functional elements on the x-axis. Dot size represents the odds-ratio of Fisher´s exact test, all tests were statistically significant (*P*-value < 0.05). First panel: Overlap with SCREEN candidate human enhancers and promoters. Second panel: Overlap with predicted promoters for selected tissues. Third panel: Overlap with predicted distal enhancers for selected tissues. Fourth panel: Overlap with predicted proximal enhancers for selected tissues.

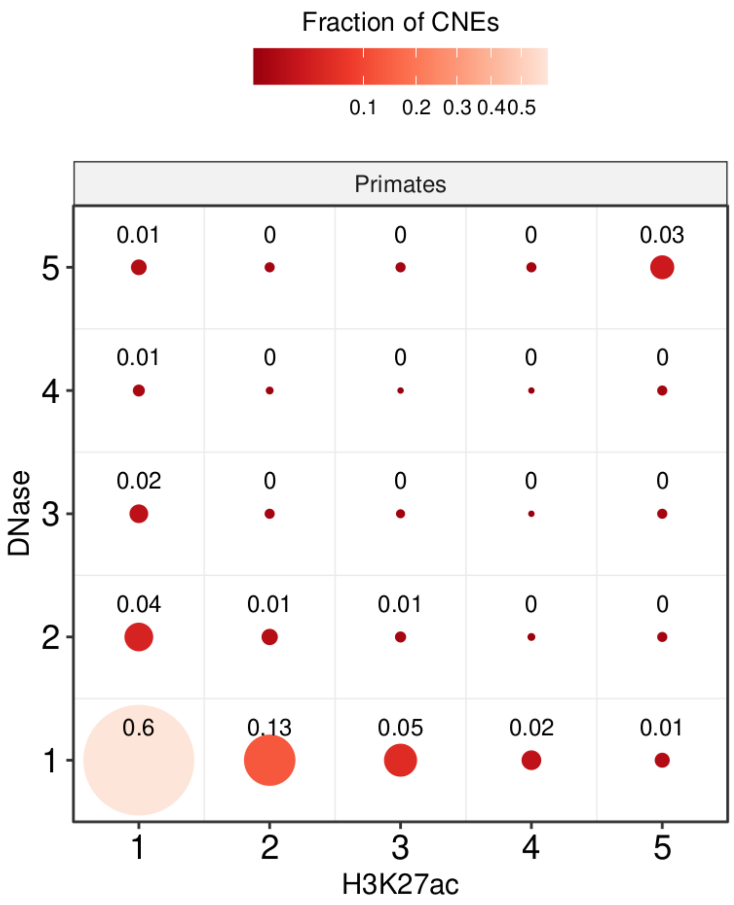


**Supplementary Figure 2. Bubble plot depicting the overlap between DNase I hypersensitivity and H3K27ac categories.** The data shows a strong positive correlation between accessibility and acetylation states.

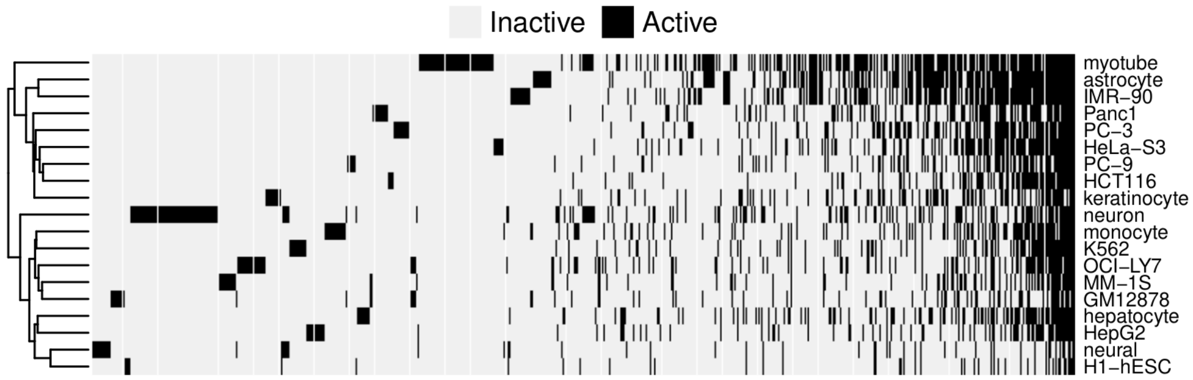


**Supplementary Figure 3. Heatmap illustrating the activity profiles of primate CNEs across 19 tissues.** Active CNEs are those in Category 4 or 5 for both DNase I hypersensitivity and H3K27ac. Only CNEs active in at least one tissue are shown. The distribution reveals a clear bimodal signature, with CNEs predominantly exhibiting either ubiquitous activity or high tissue-specificity.

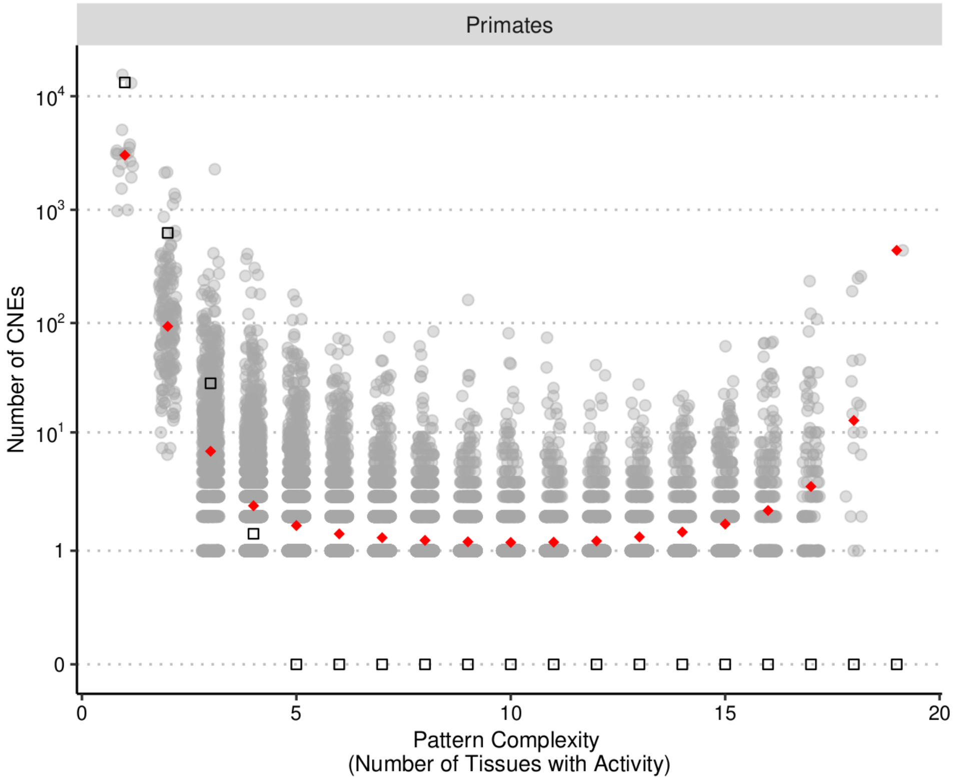


**Supplementary Figure 4. Complexity and distribution of CNE activity patterns.** Each gray dot represents an individual tissue-activity pattern. Pattern complexity is defined as the number of active tissues (*x*-axis; $n = 1,2, \ldots, 19$), with the *y*-axis showing the number of CNEs exhibiting each specific pattern for each complexity level. Red diamonds indicate the mean number of CNEs observed for each complexity level, while black circles represent the expected counts under a Binomial distribution. While the theoretical model exceeds the observed mean for low-complexity patterns ($n = 1,2,3$), the observed mean is higher for all levels where $n \geq 4$. The characteristic U-shaped distribution underscores a functional dichotomy between highly tissue-specific CNEs and those exhibiting constitutive activity across all 19 tissues.

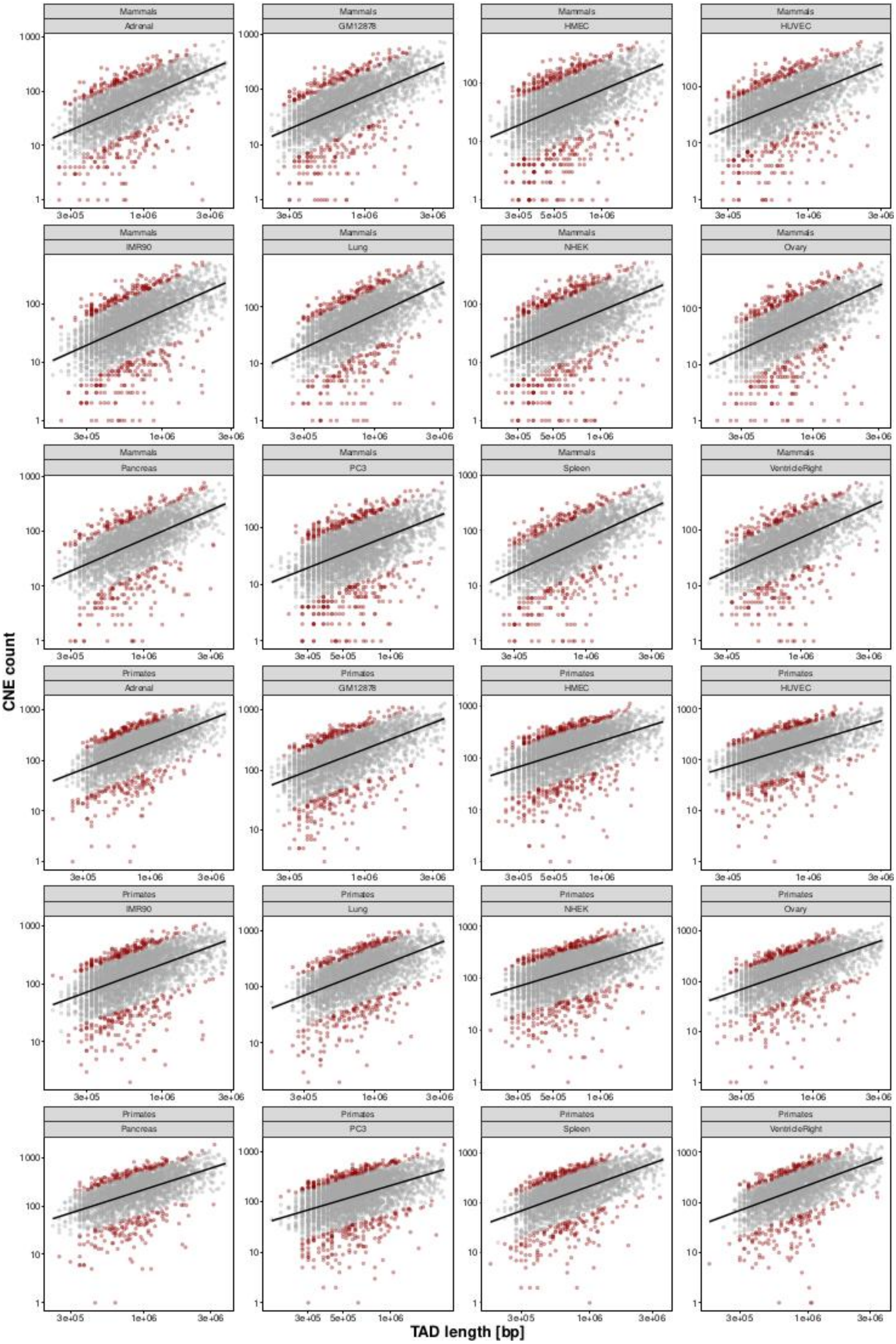
CNE count
TAD length [bp]
Mammals
Adrenal
GM12878
HMEC
HUVEC
IMR90
Lung
NHEK
Ovary
Pancreas
PC3
Spleen
VentricleRight
Primates
Adrenal
GM12878
HMEC
HUVEC
IMR90
Lung
NHEK
Ovary
Pancreas
PC3
Spleen
VentricleRight

**Supplementary Figure 5. Correlation between TAD length (in bp) and CNE density.** Scatter plots across clades and tissues illustrating the relationship between CNE number and TAD length in base pairs (bp). The number of CNEs was strongly positively correlated with TAD length. TADs identified as outliers are highlighted in dark red. Specifically, outliers were defined as the 5% of TADs with the most extreme positive and negative residuals from a linear regression of CNE count against TAD length.

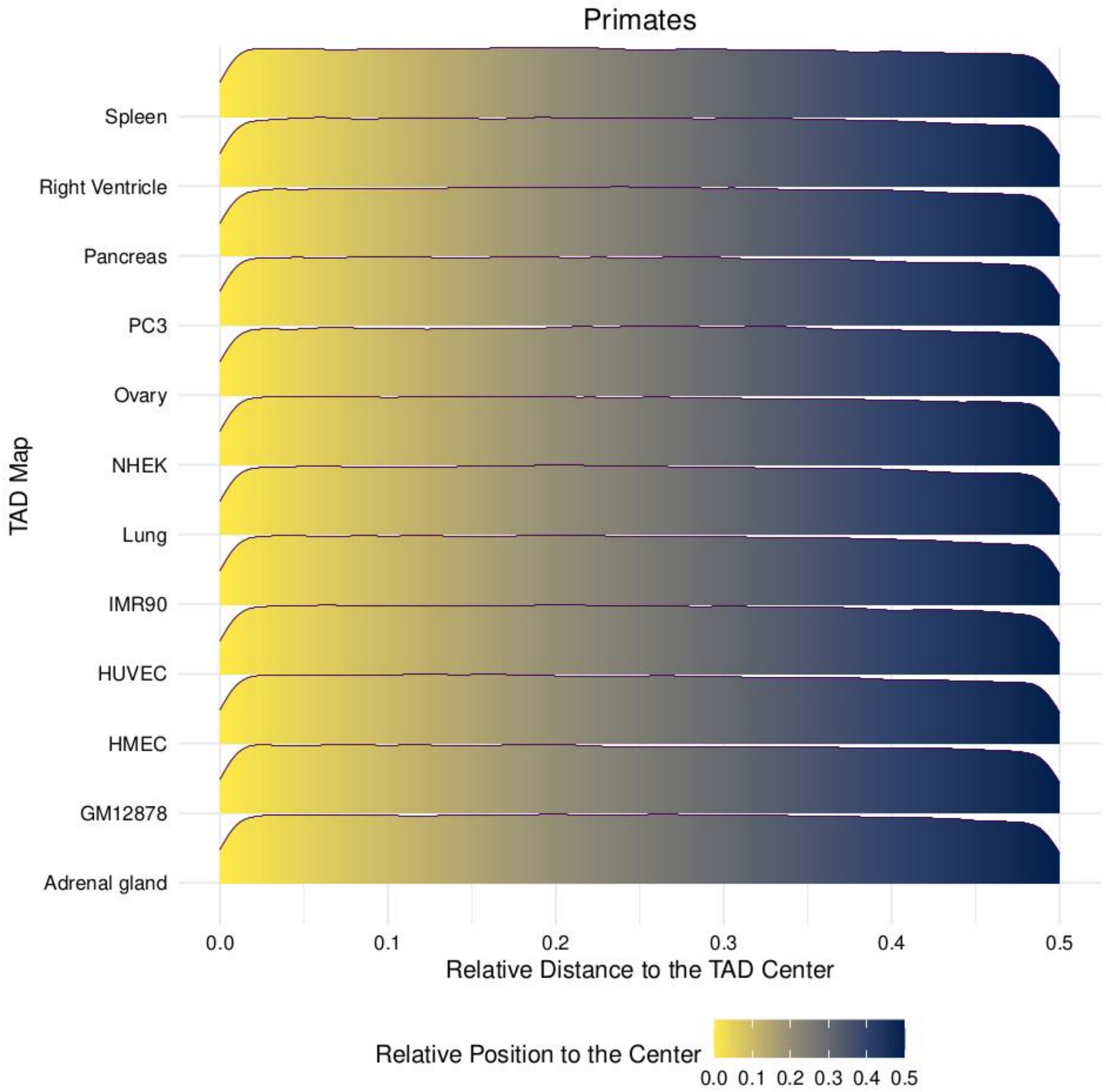


**Supplementary Figure 6. Distribution of CNEs within TADs.** Ridgeline plot showing the density distribution of primate CNEs relative to their respective TADs. CNE density peaked at TAD centers and decreased significantly toward domain boundaries.

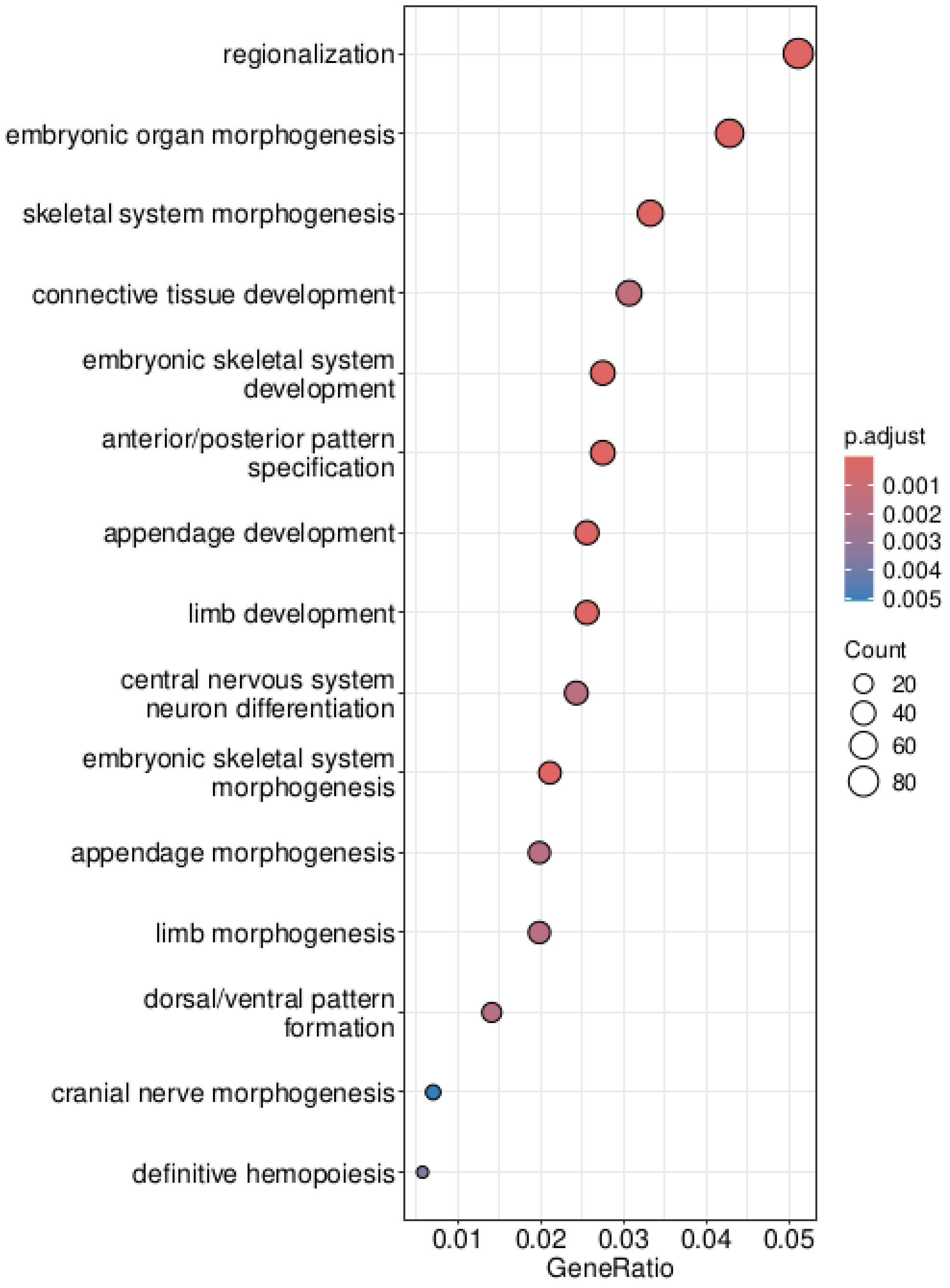


**Supplementary Figure 7. Functional enrichment analysis of primate CNE-dense TADs.** TADs with high CNE density (c) are significantly enriched for genes involved in Gene Ontology (GO) terms for development and morphogenesis. Bubble size is proportional to the number of genes associated with each GO term, while the color gradient represents the statistical significance (adjusted *P*-value).

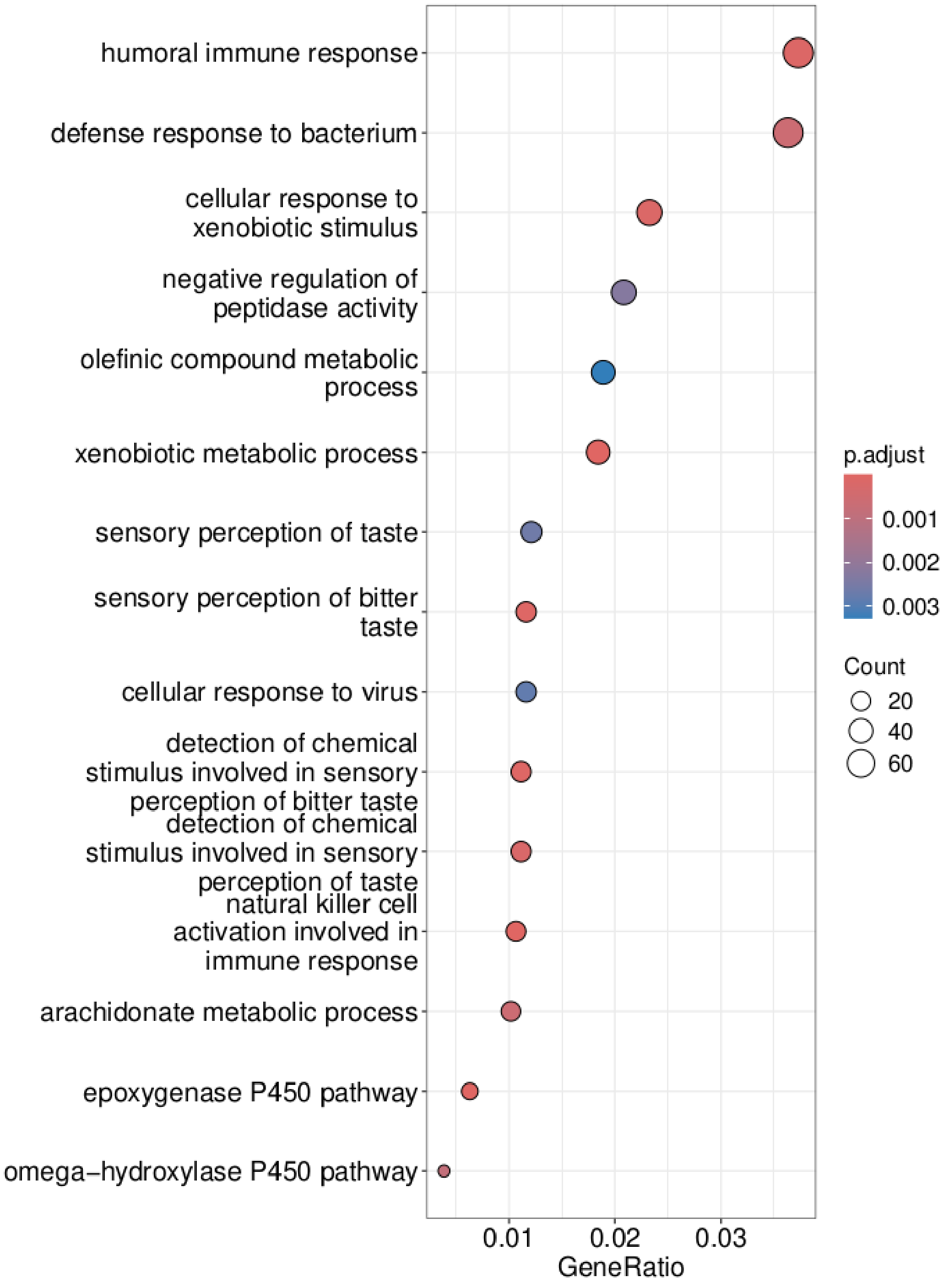


**Supplementary Figure 8. Functional enrichment analysis of primate CNE-poor TADs.** Genes in TADs with low CNE density (d) are primarily associated with responses to external stimuli and environmental factors. Bubble size is proportional to the number of genes associated with each GO term, while the color gradient represents the statistical significance (adjusted *P*-value).

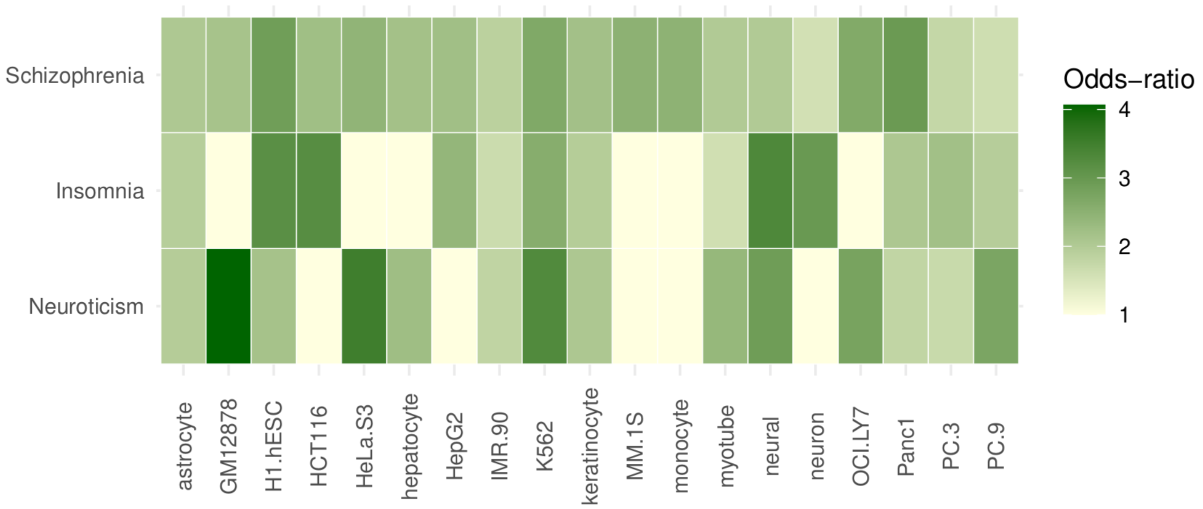


**Supplementary Figure 9. Enrichment of GWAS variants for schizophrenia, insomnia, and neuroticism within active CNEs.** Fisher's odds ratios (OR) derived from the intersection of active CNEs and trait-associated variants in the GWAS Catalog. The heatmap displays the extent to which variants for each trait are enriched among the subset of CNEs featuring activity in a particular tissue, i.e., the tissue-specific distribution of genetic burden.

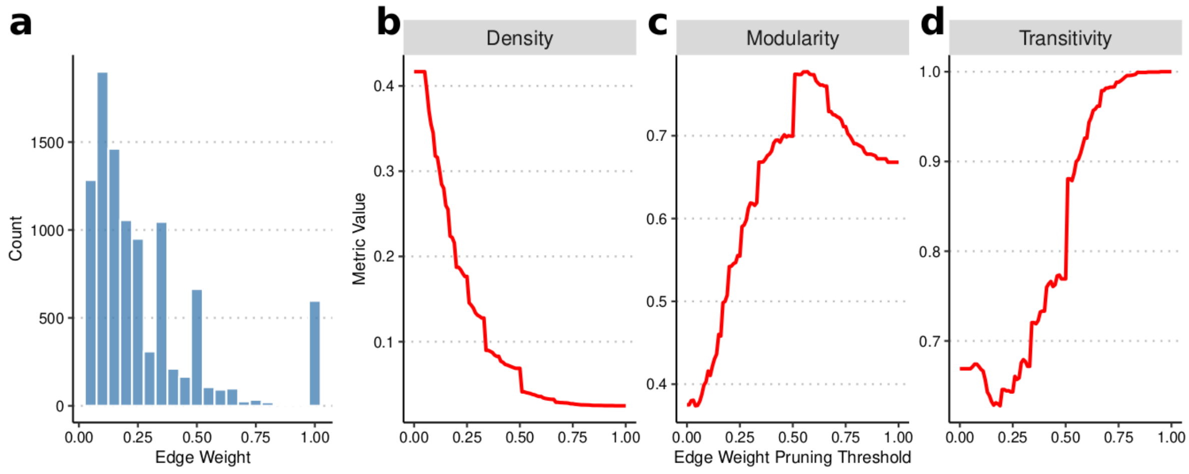


**Supplementary Figure 10. Topological changes in the CNE co-activity network upon recursive weight pruning.** (a) Distribution of edge weights in the baseline network, showing a highly skewed profile dominated by low-weight Jaccard similarity indices. (b–d) Changes in global network metrics as a function of the edge-weight pruning threshold (from low to high): (b) Network density exhibits a continuous, monotonic decrease as lower-weight edges are removed; (c) Network modularity first increases, peaking near a 50% pruning threshold before gradually declining; and (d) Network transitivity tends to decrease when excluding low-weight edges and then increases steadily toward higher pruning thresholds, indicating a highly clustered core.

### Supplementary Tables and Legends

**Supplementary Table 1. Phylogenetic distribution of CNEs.** Binary matrix representing the presence (1) or absence (NA) of CNEs as inferred through ancestral reconstruction. Columns denote the specific clade ancestor in which an element was detected. For example, a '1' in the 'Primates' column indicates conservation originating in or predating the primate common ancestor, while the full phylogenetic breadth of each CNE is defined by its cumulative presence across the matrix.

**Supplementary Table 2. Tissue-specific DNase I hypersensitivity categories of mammalian and primate CNEs.** To characterize the DNase I hypersensitivity fold-enrichment of each element, CNEs were stratified into five categories in each of the 19 tissues considered. These categories were defined by the log-transformed ratio of the average CNE DNase-seq signal relative to the mean signal across size- and chromosome-matched random regions, and range from 1 (closed chromatin) to 5 (open chromatin).

**Supplementary Table 3. Tissue-Specific H3K27ac fold-enrichment of mammalian and primate CNEs.** To characterize the acetylation of each element, CNEs were stratified into five categories (from Category 1: "no acetylation" to Category 5: "very high acetylation") based on their average H3K27ac signal (specifically, the average of log-transformed fold-change of the H3K27ac signal relative to the corresponding input control) in each of the 19 tissues considered.

**Supplementary Table 4. Tissue-Specific activity of mammalian and primate CNEs.** CNEs exhibiting an open chromatin configuration and high acetylation signal (Categories 4 and 5 for both) in a given tissue were defined as "active" *cis*-regulatory elements in that tissue. Active CNEs for a tissue are labeled with TRUE and the remaining as FALSE.

**Supplementary Table 5. Topological properties of the original CNE co-activity network.** Summary of network-wide average node-level centrality metrics calculated for the original, unpruned network with 220 nodes and 10,039 edges. Nodes represent active CNEs, with edges between any pair exhibiting a tissue-activity Jaccard similarity index greater than 0. The metrics capture different dimensions of a CNE's importance: degree quantifies a CNE's local connectivity; strength evaluates tissue-synchronization; betweenness centrality identifies critical cross-tissue regulatory bridges; and weighted PageRank distinguishes master regulators that are themselves linked to other master regulators.

| **Metric** | **Mean +/- SD** | **Median** |
|---|---|---|
| Degree | 91.264 +/- 49.884 | 85.500 |
| Strength | 24.870 +/- 11.526 | 28.632 |
| Normalized weighted betweenness | 0.005 +/- 0.009 | 0.001 |
| Weighted PageRank | 0.005 +/- 0.002 | 0.005 |

**Supplementary Table 6. Topological properties of the pruned CNE co-activity network.** Summary of network-wide average node-level centrality metrics calculated for the pruned network with 220 nodes and 2,032 edges. This topology was derived using an edge-weight threshold of 0.38, resulting in the removal of approximately 87% of low-weight edges from the original network. See Supplementary Table 5 for details.

| **Metric** | **Mean +/- SD** | **Median** |
|---|---|---|
| Degree | 18.473 +/- 12.180 | 15.000 |
| Strength | 12.198 +/- 8.668 | 9.888 |
| Normalized weighted betweenness | 0.011 +/- 0.019 | 0.003 |
| Weighted PageRank | 0.005 +/- 0.002 | 0.005 |

**Supplementary Table 7. Topological properties of the schizophrenia, insomnia and neuroticism induced subnetworks.** Summary of global network-wide properties and average node-level centrality metrics calculated for the schizophrenia, insomnia and neuroticism induced subnetworks. Nodes represent active CNEs, with edges between any pair exhibiting a tissue-activity Jaccard similarity index greater than 0.38. The observed value describes the network, the mean value together with the standard deviation describes the values across 500 independent iterations. The *P*-value describes the significance and the Benjamini-Hochberg adjusted *P*-value the significance corrected for multiple testing. The z-score shows how unusual the observed value was compared to the distribution of 100 independent iterations.

| **Trait** | **Metric** | **Observed value** | **Mean +/- SD** | ***P*-value** | **Adjusted *P*-value** | **z-score** |
|---|---|---|---|---|---|---|
| **Schizophrenia** | **Edge density** | 0.078 | 0.084 +/- 0.009 | 0.507 | 0.592 | -0.669 |
| | **Average degree** | 7.417 | 7.970 +/- 0.827 | 0.507 | 0.592 | -0.669 |
| | **Average strength** | 4.679 | 5.291 +/- 0.593 | 0.301 | 0.588 | -1.033 |
| | **Average weighted PageRank** | 0.010 | 0.010 +/- 0.000 | 1.000 | 1.000 | 0.000 |
| | **Von Neumann entropy** | 4.234 | 4.171 +/- 0.041 | 0.110 | 0.588 | 1.538 |
| | **Modularity** | 0.709 | 0.679 +/- 0.032 | 0.336 | 0.588 | 0.935 |
| **Insomnia** | **Edge density** | 0.128 | 0.085 +/- 0.012 | 0.002 | 0.004 | 3.475 |

| | | | | | | |
|---|---|---|---|---|---|---|
| | **Average degree** | 8.031 | 5.320 +/- 0.780 | 0.002 | 0.004 | 3.475 |
| | **Average strength** | 5.587 | 3.502 +/- 0.550 | 0.002 | 0.004 | 3.792 |
| | **Average weighted PageRank** | 0.016 | 0.016 +/- 0.000 | 1.000 | 1.000 | 0.000 |
| | **Von Neumann entropy** | 3.738 | 3.699 +/- 0.060 | 0.519 | 0.606 | 0.639 |
| | **Modularity** | 0.548 | 0.671 +/- 0.047 | 0.014 | 0.024 | -2.614 |
| **Neuroticism** | **Edge density** | 0.099 | 0.085 +/- 0.014 | 0.334 | 0.585 | 0.967 |
| | **Average degree** | 5.132 | 4.407 +/- 0.750 | 0.334 | 0.585 | 0.967 |
| | **Average strength** | 3.447 | 2.898 +/- 0.533 | 0.297 | 0.585 | 1.032 |
| | **Average weighted PageRank** | 0.019 | 0.019 +/- 0.000 | 1.000 | 1.000 | 0.000 |
| | **Von Neumann entropy** | 3.620 | 3.470 +/- 0.069 | 0.032 | 0.221 | 2.168 |
| | **Modularity** | 0.705 | 0.664 +/- 0.056 | 0.433 | 0.606 | 0.736 |

**Supplementary Table 8. List of 201 assemblies used for ancestral reconstruction.** Corresponding chromosome or scaffold lengths were obtained from the UCSC Genome Browser, the NCBI Genomes database, and the DNA ZOO project, as indicated in the "Source" column. For assemblies sourced from DNA ZOO, sequence lengths were calculated directly from the assembly FASTA files. Details are provided in the Methods.

| Genome Assembly | Source |
|---|---|
| hg38, panPan3, panTro6, gorGor6, ponAbe3, rheMac10, chlSab2, nasLar1, otoGar3, mm39, rn6, cavPor3, hetGla2, speTri2, ochPri3, bosTau9, balAcu1, susScr11, vicPac2, cerSim1, equCab3, neoSch1, canFam4, canFam5, felCat9, sorAra2, echTel2, triMan1, monDom5 | UCSC Genome Browser |
| HLhylMol2, HLnomLeu4, HLmacFas6, macNem1, HLpapAnu5, HLtheGel1, cerAty1, HLcerMon1, HLpilTep2, HLtraFra1, HLrhiRox2, HLsapApe1, HLcheMed1, HLgalVar2, HLmusSpr1, HLmusCar1, HLmusPah1, HLmasCou1, HLgraSur1, HLratRat7, HLacoRus1, HLacoCah1, HLpsaObe1, HLcriGri3, mesAur1, HLonyTor1, HLperEre1, HLperCal2, HLperManBai2, HLperPol1, HLmyoGla2, HLarvAmp1, HLmicFor1, micOch1, jaJac1, dipOrd2, HLereDor1, cavApe1, chiLan1, octDeg1, HLfukDam2, HLsciVul1, HLsciCar1, HLmarFla1, HLmarMar1, HLlepTim1, HLoryCunCun4, oryCun2, HLoviCan1, HLoviAmm1, HLoviAri5, HLoviOri1, HLcapSib1, HLcapIbe1, HLoreAme1, HLcapHir2, HLhipEqu1, HLsynCaf1, HLbosInd2, HLbosMut2, HLbosGru1, HLmosMos1, HLranTarGra2, HLhydIne1, HLmunRee1, HLmunMun1, HLcerHanYar1, HLaxiPor1, HLcerEla1, HLantAme1, HLtraJav1, HLdelLeu2, orcOrc1, HLlagObl1, HLgloMel1, HLsouChi1, HLturAdu2, HLturTru4, HLmonMon1, HLphoSin1, HLphyCat2, HLmegNov1, HLbalMus1, HLcatWag1, | NCBI Genomes Database |

| | |
|---|---|
| HLcamFer3, HLcamDro2, HLcamBac1, HLlamGuaCac1, HLvicVicMen1, HLdicBic1, HLequAsiAsi2, HLlonCan1, HLlutLut1, HLmusErm1, HLmusPut1, HLmusFur2, HLphoVit1, HLmirLeo1, HLcalUrs1, HLarcGaz2, HLzalCal1, HLeumJub1, HLailMel2, HLursThi1, ursMar1, HLursArc1, HLvulVul1, HLlycPic2, HLcanLupDin1, HLaciJub2, HLlynCan1, HLpumYag1, HLpumCon1, panTig1, HLpanPar1, HlpanLeo1, HLsurSur1, HLmanPen2, HLmanJav2, HLaeoCin1, eptFus1, HLpipPip2, HLstuHon1, pteAle1, HLpteGig1, HLeonSpe1, HLrouLes1, conCri1, eleEdw1, chrAsi1, HLphaCin1, HLvomUrs1, HLtriVul1, HLantFla1, HLsarHar2, HLtacAcu1, HLornAna3 | |
| HLmacFus1, HLallNig1, HLsaiBol1, HLcalPym1, HLeulFla1, HLperNas1, HLperCri1, HLcasCan3, HLconTau2, HLoryDam1, HLodoVir2, HLokaJoh2, HLgirCam2, HLpepEle1, HLeubGla1, HLequQuaBoe1, HlailFul2, enhLutKen1, HLbasSum1, HLproLot1, HLeriBar1, HLodoRos1, HLursAme2, HLlycPic3, HLneoNeb1, HLmyoSep1, HLmyoLuc1, HLeidHel2, HLeleMax1, HLchoHof3, HLphaGym1, HLmacGig1, HLospRuf1, HLnoteug3, HLdidVir1 | DNA ZOO Project Website |

**Supplementary Table 9. Candidate *enhancers and promoters* from the SCREEN database used to functionally characterize CNEs.** The reference genome assembly used for mapping is GRCh38. Database describes where the set could be found on the homepage of the SCREEN project V3. Download links were last accessed on June 11, 2026.

| Sample Label | Set | Database | Download link |
|---|---|---|---|
| GRCh38-ELS | human candidate enhancers (hg38) | Quick Start - Human | https://downloads.wenglab.org/cCREs/GRCh38-ELS.bed |
| GRCh38-PLS | human candidate promoters (hg38) | Quick Start - Human | https://downloads.wenglab.org/cCREs/GRCh38-PLS.bed |

**Supplementary Table 10. Candidate *cis*-regulatory elements (cCREs) for specific tissues from the SCREEN database used to functionally characterize CNEs.** The reference genome assembly used for mapping is GRCh38. Database describes where the set could be found on the homepage. BED files were retrieved from https://downloads.wenglab.org/Registry-V3/Seven-Group/ (last accessed on June 11, 2026).

| Sample Label | Tissue | Database | BED file |
|---|---|---|---|
| keratinocytes | skin of body - keratinocyte female donor ENCDO268AAA | Adult primary cells and tissues | ENCFF136RNO_ENCFF630BQS_ENCFF611XLA_ENCFF975BGM.7group.bed |

| | | | |
|---|---|---|---|
| OCI-LY7 | blood - OCI-LY7 | Cell lines | ENCFF414OGC_ENCFF806YEZ_ENCFF849TDM_ENCFF736UDR.7group.bed |
| K562 | bodily fluid - K562 | Cell lines | ENCFF414OGC_ENCFF806YEZ_ENCFF849TDM_ENCFF736UDR.7group.bed |
| MCF-7 | mammary gland - MCF-7 | Cell lines | ENCFF270ENA_ENCFF935BFQ_ENCFF063VLJ_ENCFF662LGI.7group.bed |
| GM12878 | bodily fluid - GM12878 | Cell lines | ENCFF428XFI_ENCFF280PUF_ENCFF469WVA_ENCFF644EEX.7group.bed |
| Panc1 | pancreas - Panc1 | Cell lines | ENCFF857RXA_ENCFF756NMQ_ENCFF493AZX_ENCFF004ITE.7group.bed |
| IMR-90 | lung - IMR-90 | Cell lines | ENCFF971HXR_ENCFF376ZIM_ENCFF699OAR_ENCFF105FHL.7group.bed |
| neural progenitor cells | neural progenitor cell originated from H9 | Adult primary cells and tissues | ENCFF286QGB_ENCFF835JIA_ENCFF618RAO_ENCFF700SCP.7group.bed |
| HepG2 | endocrine gland - HepG2 | Cell lines | ENCFF546MZK_ENCFF732PJK_ENCFF795ONN_ENCFF357NFO.7group.bed |
| monocytes | blood - CD14-positive monocyte female donor ENCDO265AAA | Adult primary cells and tissues | ENCFF389PZY_ENCFF587XGD_ENCFF184NWF_ENCFF496PSJ.7group.bed |

| | | | |
|---|---|---|---|
| myotube | musculature of body - myotube originated from skeletal muscle myoblast | Adult primary cells and tissues | ENCFF594BSS_ENCFF127SRZ_ENCFF532FVC_ENCFF450CWJ.7group.bed |
| hepatocytes | epithelium - hepatocyte originated from H9 | Adult primary cells and tissues | ENCFF902EEH_ENCFF137IUT_ENCFF347LDC_ENCFF491FMJ.7group.bed |
| A673 | musculature of body - A673 | Cell lines | ENCFF816IIS_ENCFF958CFK_ENCFF213BSP_ENCFF070LLG.7group.bed |
| HCT116 | intestine - HCT116 | Cell lines | ENCFF431JDU_ENCFF964OOU_ENCFF787LMI_ENCFF388PVO.7group.bed |
| HeLa-S3 | epithelium - HeLa-S3 | Cell lines | ENCFF757GHL_ENCFF432PYK_ENCFF658XKZ_ENCFF179RSE.7group.bed |
| astrocytes | spinal cord - astrocyte | Adult primary cells and tissues | ENCFF963PFR_ENCFF577BWJ_ENCFF643ZMC_ENCFF714NPP.7group.bed |
| MM-1S | bodily fluid - MM.1S | Cell Lines | ENCFF735XLO_ENCFF970LMB_ENCFF481LLD_ENCFF838OJW.7group.bed |
| H1-hESC | embryo - H1 | Cell lines | ENCFF573NKX_ENCFF760NUN_ENCFF919FBG_ENCFF332TNJ.7group.bed |
| neurons | brain - bipolar neuron originated from GM23338 treated with doxycycline hyclate | Adult primary cells and tissues | ENCFF386FNE_ENCFF768NPJ_ENCFF435NQW_ENCFF541XGP.7group.bed |

| | | | |
|---|---|---|---|
| PC-9 | lung - PC-9 | Cell lines | ENCFF623TJB_ENCFF465MDM_ENCFF907RYE_ENCFF936QRH.7group.bed |
| PC-3 | prostate gland - PC-3 | Cell lines | ENCFF599UKS_ENCFF319OET_ENCFF537PUA_ENCFF756ESH.7group.bed |

**Supplementary Table 11. DNase-seq datasets.** List of tissues, ENCODE accessions, and originating laboratories. Primary alignment files (BAM format) were retrieved from the ENCODE Portal using the listed accessions.

| **Sample Label** | **ENCODE ID** | **Laboratory** | **Biosample** |
|---|---|---|---|
| keratinocytes | ENCFF111DMK | John Stamatoyannopoulos, UW | keratinocyte female donor ENCDO268AAA |
| OCI-LY7 | ENCFF770HOF | John Stamatoyannopoulos, UW | OCI-LY7 |
| K562 | ENCFF156LGK | John Stamatoyannopoulos, UW | K562 |
| GM12878 | ENCFF658WKQ | John Stamatoyannopoulos, UW | GM12878 |
| Panc1 | ENCFF048ENW | John Stamatoyannopoulos, UW | Panc1 |
| IMR-90 | ENCFF023JFG | John Stamatoyannopoulos, UW | IMR-90 (female embryo 16 weeks) |
| neural progenitor cells | ENCFF048NZA | John Stamatoyannopoulos, UW | neural progenitor cell originated from H9 |
| HepG2 | ENCFF097NAZ | John Stamatoyannopoulos, UW | HepG2 |

| | | | |
|---|---|---|---|
| monocytes | ENCFF560DOD | John Stamatoyannopoulos, UW | CD14-positive monocyte female donor ENCDO265AAA |
| myotube | ENCFF173TWL | John Stamatoyannopoulos, UW | myotube (originated from skeletal muscle myoblast) |
| hepatocytes | ENCFF761VCZ | John Stamatoyannopoulos, UW | hepatocyte originated from H9 |
| A673 | ENCFF348KWA | John Stamatoyannopoulos, UW | A673 |
| HCT116 | ENCFF391EDU | John Stamatoyannopoulos, UW | HCT116 |
| HeLa-S3 | ENCFF912JKA | John Stamatoyannopoulos, UW | HeLa-S3 |
| astrocytes | ENCFF586NXB | John Stamatoyannopoulos, UW | astrocyte |
| MM-1S | ENCFF327EAE | John Stamatoyannopoulos, UW | MM.1S |
| H1-hESC | ENCFF546PJU | John Stamatoyannopoulos, UW | H1(-hESC) |
| neurons | ENCFF297XNU | John Stamatoyannopoulos, UW | bipolar neuron_originated_from_GM23338_treated_with_doxycycline_hyclate |
| PC-9 | ENCFF107HZD | John Stamatoyannopoulos, UW | PC-9 |
| PC-3 | ENCFF743EZY | John Stamatoyannopoulos, UW | PC-3 |

**Supplementary Table 12. H3K27ac ChIP-seq data sets.** List of tissues, ENCODE accessions, and originating laboratories. Primary alignment files (BAM format) were retrieved from the ENCODE Portal using the listed accessions.

| **Sample Label** | **ENCODE ID Replicate 1** | **ENCODE ID Replicate 2** | **ENCODE ID Control** | **Laboratory** | **Biosample** |
|---|---|---|---|---|---|
| keratinocytes | ENCFF578IKI | ENCFF828BCB | ENCFF292GHV | Bradley Bernstein, Broad | keratinocyte_female_donor_ENCDO268AAA |
| OCI-LY7 | ENCFF204UGN | ENCFF489RAL | ENCFF046QZN | Bradley Bernstein, Broad | OCI-LY7 |
| K562 | ENCFF301TVL | ENCFF879BWC | ENCFF392XRJ | Bradley Bernstein, Broad | K562 |
| GM12878 | ENCFF804NCH | ENCFF948GTC | ENCFF651UIO | Bradley Bernstein, Broad | GM12878 |
| Panc1 | ENCFF960KET | ENCFF384KMQ | ENCFF675MQQ | Peggy Farnham, USC | Panc1 |
| IMR-90 | ENCFF146UYU | ENCFF071VOI | ENCFF465UIJ | Bing Ren, UCSD | IMR-90(_female_embryo_16_weeks) |
| neural progenitor cells | ENCFF802FBW | ENCFF826AZF | ENCFF175BOB | Bradley Bernstein, Broad | neural_progenitor_cell_originated_from_H9 |
| HepG2 | ENCFF805KGN | ENCFF686HFQ | ENCFF645CCA | Bradley Bernstein, Broad | HepG2 |
| monocyte | ENCFF835ZFV | ENCFF329AXT | ENCFF075YNC | Bradley Bernstein, Broad | CD14-positive_monocyte_female_donor_ENCDO265AAA |

| | | | | | |
|---|---|---|---|---|---|
| myotube | ENCFF302FHM | ENCFF198UAJ | ENCFF408COK | Bradley Bernstein, Broad | myotube( originated from skeletal muscle myoblast) |
| hepatocytes | ENCFF891WDF | ENCFF098YME | ENCFF248XON | Bradley Bernstein, Broad | hepatocyte_originated_from_H9 |
| A673 | ENCFF737FZT | ENCFF918ACD | ENCFF170UTG | Bradley Bernstein, Broad | A673 |
| HCT116 | ENCFF689GKI | ENCFF340TPS | ENCFF064LQH | Bradley Bernstein, Broad | HCT116 |
| HeLa-S3 | ENCFF113QJM | ENCFF218BUW | ENCFF432JPE | Bradley Bernstein, Broad | HeLa-S3 |
| astrocytes | ENCFF084MDC | ENCFF496OLR | ENCFF216TIF | Bradley Bernstein, Broad | astrocyte |
| MM-1S | ENCFF605DAC | ENCFF937ZTU | ENCFF550VRT | Bradley Bernstein, Broad | MM.1S |
| H1-hESC | ENCFF238SQN | ENCFF242PAC | ENCFF064DDT | Bradley Bernstein, Broad | H1(-hESC) |
| neurons | ENCFF017WUP | ENCFF751YAL | ENCFF687LIL | Bradley Bernstein, Broad | bipolar_neuron_originated_from_GM23338_treated_with_doxycycline_hyclate |
| PC-9 | ENCFF655AIP | ENCFF237EXU | ENCFF754MHP | Bradley Bernstein, Broad | PC-9 |
| PC-3 | ENCFF474QFK | ENCFF174JIG | ENCFF607ZQL | Bradley Bernstein, Broad | PC-3 |

**Supplementary Table 13. TAD data sets.** List of tissues, and file names. TAD files (BED format) were retrieved from the 3D Genome Browser.

| **Sample Label** | **Downloaded file** |
|---|---|
| Ventricle Right | Right_Ventricle_GSE87112_tad.bed |
| Spleen | Spleen_GSE87112_tad.bed |
| Pancreas | Pancreas_GSE87112_tad.bed |
| Ovary | Ovary_GSE87112_tad.bed |
| NHEK | NHEK_GSE63525_tad.bed |
| Lung | Lung_GSE87112_tad.bed |
| PC3 | PC3_GSE172099_tad.bed |
| IMR90 | IMR-90_GSE63525_tad.bed |
| HUVEC | HUVEC_GSE63525_tad.bed |
| HMEC | HMEC_GSE63525_tad.bed |
| GM12878 | GM12878_GSE63525_tad.bed |
| Adrenal Gland | Adrenal_Gland_GSE87112_tad.bed |
